# A flexible and generalizable model of online latent-state learning

**DOI:** 10.1101/443234

**Authors:** Amy L Cochran, Josh M Cisler

## Abstract

Many models of classical conditioning fail to describe important phenomena, notably the rapid return of fear after extinction. To address this shortfall, evidence converged on the idea that learning agents rely on latent-state inferences, i.e. an ability to index disparate associations from cues to rewards (or penalties) and infer which index (i.e. latent state) is presently active. Our goal was to develop a model of latent-state inferences that uses latent states to predict rewards from cues efficiently and that can describe behavior in a diverse set of experiments. The resulting model combines a Rescorla-Wagner rule, for which updates to associations are proportional to prediction error, with an approximate Bayesian rule, for which beliefs in latent states are proportional to prior beliefs and an approximate likelihood based on current associations. In simulation, we demonstrate the model’s ability to reproduce learning effects both famously explained and not explained by the Rescorla-Wagner model, including rapid return of fear after extinction, the Hall-Pearce effect, partial reinforcement extinction effect, backwards blocking, and memory modification. Lastly, we derive our model as an online algorithm to maximum likelihood estimation, demonstrating it is an efficient approach to outcome prediction. Establishing such a framework is a key step towards quantifying normative and pathological ranges of latent-state inferences in various contexts.

**Author summary:** Computational researchers are increasingly interested in a structured form of learning known as latent-state inferences. Latent-state inferences is a type of learning that involves categorizing, generalizing, and recalling disparate associations between observations in one’s environment and is used in situations when the correct association is latent or unknown. This type of learning has been used to explain overgeneralization of a fear memory and the cognitive role of certain brain regions important to cognitive neuroscience and psychiatry. Accordingly, latent-state inferences are an important area of inquiry. Through simulation and theory, we establish a new model of latent-state inferences. Moving forward, we aim to use this framework to measure latent-state inferences in healthy and psychiatric populations.

## Introduction

Learning and decision-making are fundamental aspects of day-to-day human life. Indeed, many mental health disorders can be conceptualized from the perspective of biases or errors in learning and decision-making [1]. Accordingly, the study of how humans learn and make decisions is an important topic of inquiry and the past two decades has witnessed a significant surge in the application of computational modeling approaches to the problem of human learning and decision-making [2–5]. One significant insight this literature has made is the differentiation of model-free and model-based learning [6, 7]. Model-free learning refers to relatively simple updating of cached values based on trial-and-error experience. An early and influential example of model free learning is the Rescorla-Wagner (RW) model [8], which proposed that associative strength (i.e. the degree to which a cue predicted an outcome) updates in response to new experiences in proportion to the magnitude of a prediction error (i.e., how wrong the current prediction was). The RW model, and similar model-free formulations that it inspired [9, 10], powerfully explained many aspects of learning.

Nonetheless, a notable problem for model-free accounts of learning was the phenomenon of rapid recovery of responding following extinction learning [11]. That is, model-free accounts of learning based on trial-by-trial updating predict that the return of responding following extinction should essentially occur at the same rate as initial learning. This prediction is not supported by a wealth of data demonstrating that extinguished responding can rapidly return via reinstatement (i.e., an un-signalled presentation of the outcome), context renewal (i.e., returning to the initial acquisition learning context), or spontaneous recovery (i.e., return of responding following the passage of time) [12]. This rapid return of extinguished responding is an important phenomenon with implications for treatment of clinical disorders such as anxiety disorders and substance use [13, 14].

One of the first applications of a model-based account of learning that could address rapid renewal of responding following extinction was by Redish and colleagues in 2007 [11]. They argued that an agent performs two concurrent tasks during a learning experiment: 1) updating associative strength between a cue and an outcome, and 2) recognizing the current state of the experiment and storing separate associative strengths for the acquisition and extinction contexts. Rapid recovery of responding can be explained by the agent inferring that the current state of the experiment has changed to the acquisition phase and therefore responding to the cue with the associative strength stored for the acquisition phase. Though this initial formulation had limitations [15], it stimulated subsequent research developing latent-state models of reinforcement learning [16–19]. For example, if an agent is assumed to infer latent states to explain observed relationships between stimuli, actions, and outcomes, and if inferring separate latent states for acquisition and extinction phases of an experiment explains rapid recovery of responding, then it follows that blurring the experimental distinction between acquisition and extinction phases would result in less recovery of responding following extinction. That is, if the contingency between cue and outcome (e.g., 50% contingency) during the acquisition phase slowly transitions to extinction, rather than an abrupt change, then an agent is less likely to infer a separate extinction context and more likely to modify the initial acquisition memory. Because the acquisition associative strength is lower, less subsequent recovery of responding would be expected in a recall test. A carefully designed experiment using an animal model demonstrated exactly this observation, providing strong evidence for a latent-state model of learning [20].

The application and utility of latent-state learning models is not confined to explaining recovery of responding following extinction. Tolman’s early theory of cognitive maps [21] posits that an agent forms abstract representations of a task’s state space. There has been a recent resurgence of interest in this concept of cognitive maps in the cognitive neuroscience field informed by latent-state computational models [17, 22, 23]. This work, conducted both using animal and human experimental models, suggests that essential functions of the orbitofrontal cortex (OFC) and hippocampus are the encoding and representation of latent task space (i.e., forming cognitive maps of a task’s state space) [23]. This theory potentially powerfully describes a superordinate process that explains why the OFC and hippocampus are implicated in such a diverse array of cognitive and emotional functions. This theory also has implications that tie back into extinction learning and clinical disorders and suggest a novel understanding of the role of the OFC in mediating poor extinction learning. However, it is important to recognize that the role of the OFC remains highly debated.

As can be seen, latent-state theories have significant implications for our understanding of normative and disordered mechanisms of learning. The purpose of the current work is to introduce a new model of latent-state learning and to verify the suitability and utility of this model for explaining group-level effects of classical conditioning. The introduced model makes six key assumptions about how an agent performs latent-state inferences, notably that a learning agent uses latent states to index disparate *associations* between cues in an effort to predict rewards and that they assume latent states are relatively stable over time. Altogether, these assumptions differentiate our model from past efforts to describe latent state learning [11, 15, 17, 19]. These differences are highlighted in a series of simulation experiments. In addition to formalizing a new theory of latent-state learning, we were also motivated to provide the research community with a practical model that can be fit to data from a diverse set of experiments probing classical conditioning and latent-state learning. Even though every model has limitations and cannot be applied to every experiment, there is practical significance in being able to compare fitted parameters from a model across experiments.

## Results

### Model

We present a model of how an agent uses latent states to learn psychological experiments consisting of a sequence of trials, wherein each trial a learning agent is exposed to a collection of cues followed by rewards (and/or penalties). We use the following notation. Cue *n* on trial *t* is denoted by *c_n_*(*t*) and takes a value of 1 when the cue is present and 0 otherwise. Cues are collected in a vector 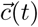. Rewards on trial *t* are denoted by *R*(*t*). After trial *t*, the strength of the association between cue *n* and rewards is denoted by *V_n_*(*t*). An apostrophe denotes transposition, and the notation *X*(1:*t*) is shorthand for (*X*(1)*, …, X*(*t*)).

Our model assumes a learning agent has a certain world view about how rewards are generated and wants to predict rewards efficiently based on this world view (Fig 1A). Their world view assumes rewards are a function of which cues are presented, a latent state, and a latent error. A latent state is an index to disparate associations between cues and rewards. In order to predict rewards, they must invert their view of the world to infer which latent state is active, how cues are associated with rewards for each latent state, and the expected uncertainty in rewards due to the latent error. We proceed to describe our model for how an agent performs these tasks. We leave details to the Methods section about how our model of latent-state learning can be derived formally from this world view.

**Fig 1.**
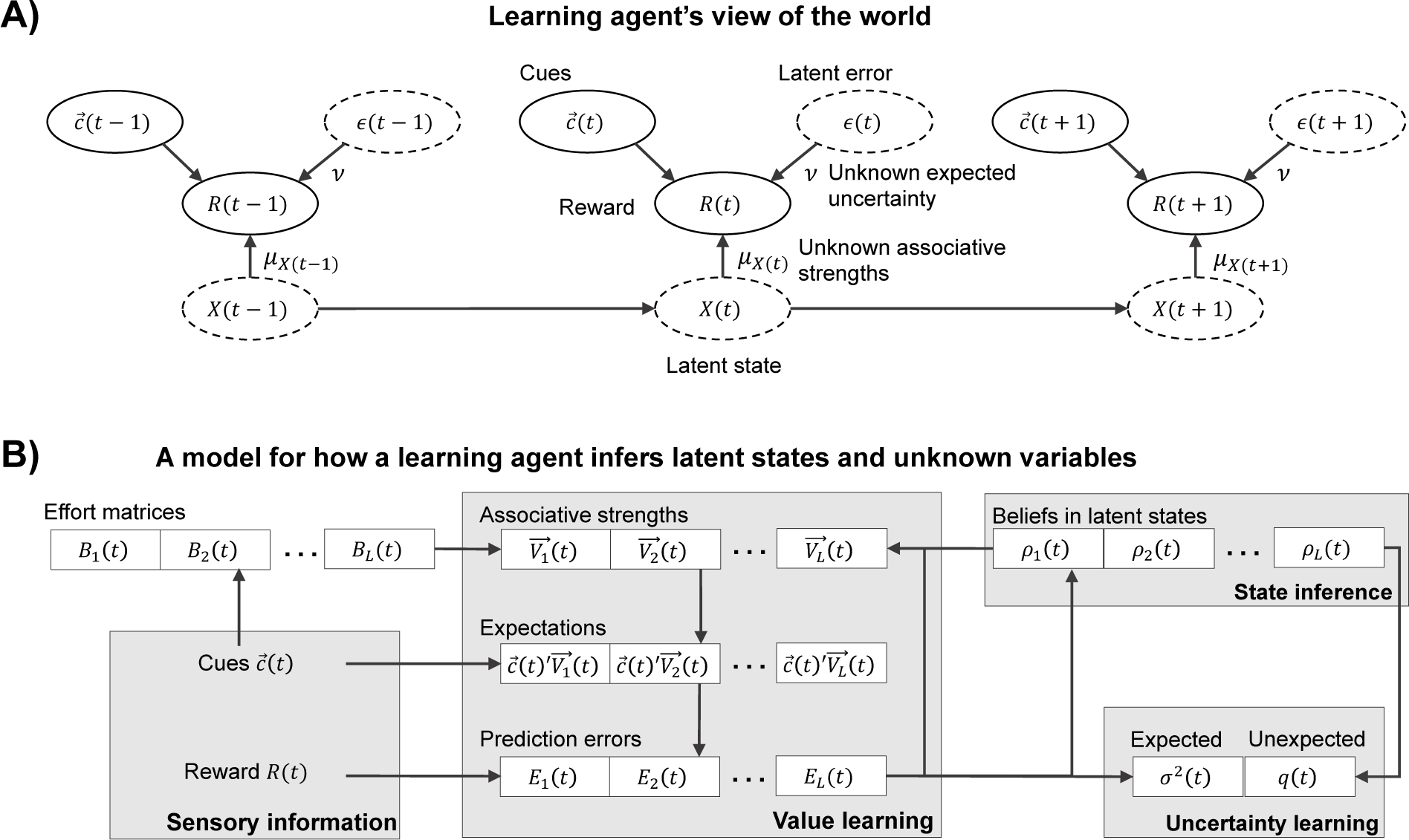
A) A learning agent’s world view whereby rewards are generated according to cues, a latent state, and a latent error. In order to predict rewards, they must infer which latent state is active, the relationship between cues and rewards for each latent state, and the expected uncertainty in rewards due to the latent error. B) The proposed model for how a learning agent inverts their world view. They first observe cues to generate expectations or predictions for rewards based on *L* estimates of associative strengths corresponding to *L* latent states. Upon observing rewards, they use errors in their predictions to update associative strengths, measures of uncertainty, and beliefs in which state is active. The degree to which associative strengths can be updated depends on both the agent’s belief in the corresponding latent state and the corresponding effort matrix, which keeps track of how cues covary.

#### Building upon the Rescorla-Wagner (RW) model

If an agent believed in only one latent state, then a RW model could be used to determine the relationship between cues and rewards. The RW model proposes a linear relationship between cues and expected rewards.

This relationship is captured by an associative strength for each cue updated according to:

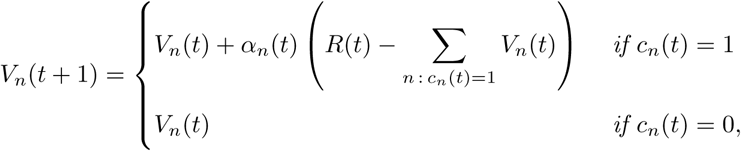

with initial associative strength *V_n_*(1) = 0. Term *α_n_*(*t*) is referred to as *associability* and is constant in the RW model. Collecting associative strengths in a vector 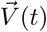, we can express this update concisely as

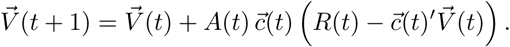

The term *A*(*t*) is now a matrix capturing associability. The RW model assumes *A*(*t*) is diagonal with terms *α_n_*(*t*) along its diagonal, whereas we will consider matrices that are not diagonal. Changes in associative strength depends on the learning agent’s current expectation or best guess 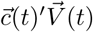 for rewards given observed cues. If their expectation exceeds the actual reward, then associative strength declines for each cue present. If their expectation is below the actual reward, then associative strength increases for each cue present. By updating associative strength in vector form, we see that rewards are captured by a linear regression model of cues. Hence, cues can be continuous or represent interactions, i.e. the presence or absence of a compound of cues. That is, the same update formula applies for continuous cues and interaction terms. Also note that the RW model learns cue-reward associations rather than action-value associations as in temporal difference reinforcement learning.

#### Allowing for latent-state learning

We build upon the RW model by allowing a learning agent to propose *L* competing (i.e. mutually-exclusive) *associations* between cues and rewards. The index to each association is referred to as a *latent state* and is associated with its own RW model for predicting rewards from cues. The agent learns about associations for each latent state while learning about their belief in which latent state best explains current observations. We capture belief in latent state *l* on trial *t* as a positive variable *p_l_*(*t*) such that beliefs sum to one: *p*_1_(*t*) + *…* + *p_L_*(*t*) = 1.

We let 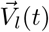 represent the vector of associative strengths and *A_l_*(*t*) represent the associability matrix for latent state *l*. The learning agent updates associative strength as before except for the subscript *l*:

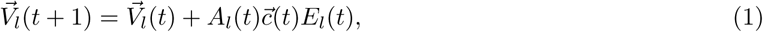

where we used the shortened notation *E_l_*(*t*) to represent the prediction error for latent state *l* on trial *t*:

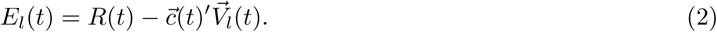

#### An associability process

Associability is constant in the RW model. By contrast, experiments such as those by Hall and Pearce [9, 24] suggest associative strength is path-dependent: associability, i.e. how quickly associative strength changes, depends on the history of observations and changes over time [10]. We describe associability as depending on current beliefs and a matrix *B_l_*(*t*) which we refer to as an *effort* matrix:

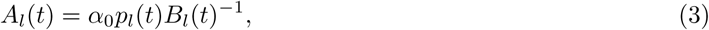

for a patent-specific parameter *α*_0_ *∈* [0, 1] controlling how much an individual weights older observations versus newer observations. The effort matrix *B_l_*(*t*) is updated according to

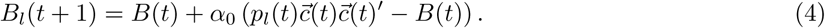

with *B_l_*(1) = *I*. The effort matrix estimates the matrix of cue second moments associated with each latent state. Second moment information of cues has been integrated into other models of learning, but usually in the form of variances and covariances rather than second moments directly [11, 19]. The use of the effort matrix is motivated in our derivation of our model in the Methods section and is similar to matrices found in an online algorithm for solving the machine learning problem of contextual bandits with linear rewards [25].

#### Updating latent-state beliefs

Beliefs link latent states in our model. The assumed world view of the learning agent proposes that rewards are generated on trial *t* as follows: a latent random variable *X*(*t*) *∈ {*1*, …, L}* is drawn from some distribution and rewards are drawn depending on the current latent state *X*(*t*) and cues 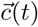 (Fig 1A). Further, rewards are assumed to be mutually independent between trials conditional on latent states, and latent states are assumed to be Markovian, i.e. *X*(*t*) depends on *X*(1:*t −* 1) only through *X*(*t −* 1). Under these assumptions, Bayes law yields a posterior distribution over latent states from observed rewards given by:

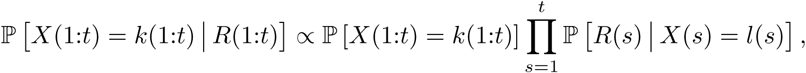

where we suppress the dependence on cue vectors to shorten notation. The posterior probability *ρ_l_*(*t*) that latent state *X*(*t*) is *l* based on observations up to trial *t* is then:

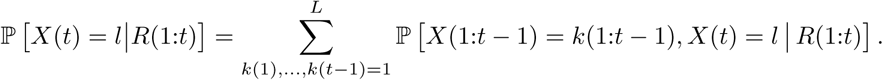

From this expression, we see that optimal Bayesian inference requires enumerating over all possible sequences of latent states, the number of which grows exponentially with *t*. Such computation is generally considered optimistic for human computation [26]. Efforts are made to reduce computation such as with particle methods [17], assuming one transition between states [27], or using a functional form of memory that does not maintain the entire history of observations.

We capture a more general dynamic learning environment similar to [26], by assuming latent states are a Markov chain in which a latent variable transitions to itself from one trial to the next with probability 1 *− γ*(*L −* 1)*/L* and transitions to a new state (all new states being equally-likely) with probability *γ*(*L −* 1)*/L*. This assumption reflects that learning often involve blocks of consecutive trials, or stages, in which rewards are generated in an identical manner. An optimal Bayesian filtering equation could then be used to update beliefs

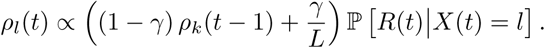

Unfortunately, this equation requires the learning agent knows the probability distribution of rewards for each latent state. We thus do not think the agent reasons in an optimal Bayesian way. Instead, we initialize beliefs *p_l_*(0) = 1*/L* and propose that the agent uses an *approximate Bayesian filtering equation* (cf. [28])

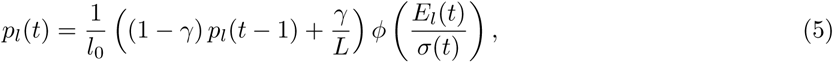

where *ϕ* is the probability density function of a standard normal random variable; *σ*(*t*) is the agent’s estimate of the standard deviation of rewards at the start of trial *t*; and

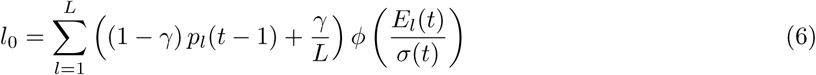

is a normalizing constant to ensure beliefs sum to 1. In other words, we replace the distribution/density function of rewards in the optimal Bayesian filtering equation with a normal density function with mean given by the agent’s current estimate 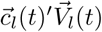 and standard deviation given by the agent’s current estimate *σ*(*t*). Even though we use a normal density function, rewards do not need to be normally-distributed or even continuous.

#### Measuring expected uncertainty

The learning agent may also track the standard deviation *σ*(*t*) of prediction errors. With only one standard deviation, we assume the learning agent estimates the standard deviation *pooled* over each latent state. We use pooled estimates to reduce the number of variables, but it may be more realistic to use a separate estimate for each latent state. We use the following update:

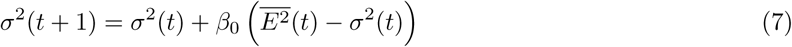

where 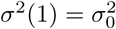 is some initial estimate of the variance; *β*_0_ is a constant associability parameter for variance *σ*^2^(*t*); and 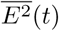 is the squared prediction error averaged over latent states:

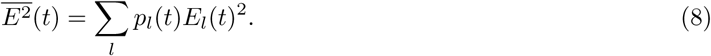

Variance is also used in other models of latent-state learning, such as the model in Redish et al [11] and the model in Gershman et al [19]. Further, Yu and Dayan [26] proposed a neural mechanism, via acetylcholine signalling, by which a learning agent keeps track of expected uncertainty in changing environments. As a staple to statistical inference, variance is a natural candidate for quantifying expected uncertainty.

#### Measuring unexpected uncertainty

Many latent-state learning models assume the number of latent state can grow [11, 15, 17]. Similarly, we use an *online* mechanism to trigger an addition of new latent states when needed to explain rewards. This mechanism reflects the need to capture *unexpected uncertainty*, as discussed in Yu an Dayan [26]. It is based on Page’s algorithm [29] for solving a problem known as change point detection (cf. [30]). The idea is to test whether we can reject our current model in favor of an alternative model with an extra latent state.

Specifically, we let *L* be the number of latent states actively considered on trial *t*. We keep track of model performance using a statistic *q*(*t*):

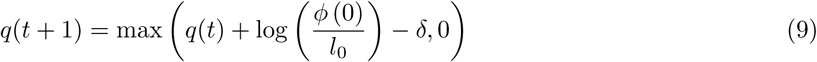

with a scalar parameter *δ* and initialized *q*(1) = 0. Importantly, *ϕ* (0) */l*_0_ is the likelihood ratio between a one-state model with expected rewards given by actual rewards *R*(*t*) and the current model, where *l*_0_ was the normalizing constant for beliefs at Eq (6). If *q*(*t*) exceeds a threshold *η*, we let

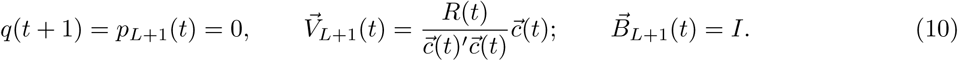

Note that 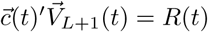 so that the current reward is the expected reward for the new state. We then replace *L* with *L* + 1.

#### Changes in context

Learning often involves changes in context whether it be new surroundings, backdrop, or point in time. Context changes are believed to influence learning in a distinctly different way than a cue influences learning [12]. Our model supposes that these changes in context corrodes current beliefs, whereby beliefs in a latent state on a previous trial are thought to be less informative on the present trial. For a temporal shift in context, our model supposes this corrosion increases with time. While the Gershman (2017) model uses a temporal kernel, we replace beliefs *p_l_*(*t*) at the end of trial *t* with

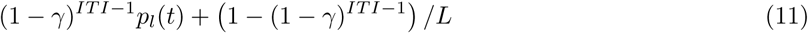

where *ITI* is the time between trials within a task phase. This update amounts to repeatedly updating latent state beliefs for *ITI −* 1 iterations according to the Markov chain transition probabilities. Beliefs would become evenly distributed between latent states in the limit as *ITI* goes to infinity. For a visual or spatial shift in context, we set *ITI* = *∞* in equation above, leading to uniform beliefs 1*/L*, i.e. all latent states are believed to be equally likely. As a result, an agent does not carry their beliefs forward to the next trial when there is a spatial or visual change of context.

A temporal shift in context might also alter how associative strengths and beliefs are updated. Gershman et al [19] describes the influence of retrieval on a memory by way of “rumination”. This mechanism is captured in the model of Gershman (2017) [19] and in our model by repeatedly replacing associative strengths 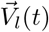 with the new estimate 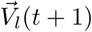 performing the updates at the end of trial *t* again. This process is repeated for a number of trials given by min*{χ, ITI −* 1*}* where *χ* is a patient-specific parameter.

#### Putting it all together

Our latent-state model of learning is summarized in Fig 1B and in Algorithm 1. Briefly, a learning agent undergoes the following each trial: they observe cues, generate expectations for rewards, observe a reward, measure error, update their belief in latent states, and update associative strength and measures of uncertainty. In between trials, a learning agent may adjust their beliefs and associative strengths to account for contextual shifts whether they be visual, spatial, or temporal. Further, parameters can be tuned to describe differences in learning between agents: *α*_0_ influences the rate of learning associative strengths, *β*_0_ influences the rate of learning variance, *γ* influences transitions between latent state, *σ*_0_ influences initial expected uncertainty, *ν* and *δ* influence unexpected uncertainty, and *χ* determines rumination.

Our model is grounded by six major predictions or assumptions about how an agent learns:

**1) An agent learns *online*.** Model variables are updated on each trial using only current observations and current values of model variables. Consequently, an agent can learn without remembering observations of cues and rewards from prior trials. Their memory and computational requirements grow only in the number of latent states rather than the number of trials, which may be a more realistic reflection of how humans learn. The Rescorla-Wagner model and other models also use online updates for certain model variables [8, 11, 17, 19].
**2) An agent uses latent states to *predict rewards*.** In deriving our model (see Methods section), we start with the assumption that the agent’s goal is to predict rewards. The role of a latent state is to account for different predictions that can be made upon observing cues. By contrast, the role of a latent state could be to discriminate between different sets of cues-reward observations [11, 17], which in turn could be used to predict rewards.
**3) Associability, i.e. how quickly associations are updated, depends on beliefs in latent states and cue novelty.** Associative strength updates quickly for latent states believed to be active, but slowly for latent states believed to be inactive. This feature is similar to the Mackintosh model [31] in that associability increases when rewards are accurately predicted. However, associative strength also updates quickly when an agent switches their belief to a new latent state to account for unexpected or surprising outcomes, i.e. outcomes in which prediction error is larger relative to the expected uncertainty. This feature is similar to Pearce-Hall model [9], which supposes associability increases in the presence of uncertainty. Meanwhile, the effort matrix, which tracks how often a cue appears (diagonal entries) and a pair of cues appear (off-diagonal entries), also determines associability. As a result, associative strength is updated more quickly when novel sets of cues are presented.
**4) Latent states are relatively stable between trials.** Latent states are expected to be the same from one trial to the next with a certain probability. This assumption causes beliefs in latent states to be relatively consistent between trials. An alternative is to assume latent states are exchangeable between trials, allowing beliefs in latent states to shift more sharply between trials [17]. Stability between trials, however, might degrade with time. Our model assumes that with time, the last trial becomes less and less informative about the latent state, to the point that the agent eventually believes each latent state is equally likely regardless of prior beliefs.
**5) Beliefs are maintained over multiple latent states.** Our model allows an agent to maintain beliefs over multiple latent states on each trial as opposed to believing in only latent state. For example, an agent is able to believe that two latent states are equally likely. This allows the agent to explicitly state their beliefs in competing associations, as is required by some learning tasks [32]. Maintaining beliefs over multiple states is a common feature for Bayesian models [17, 19, 27, 32]. An alternative is to classify each trial to one state [11, 33].

##### Algorithm 1

Proposed model of latent-state learning which updates variables *online*, i.e. does not keep track of past cues and rewards.

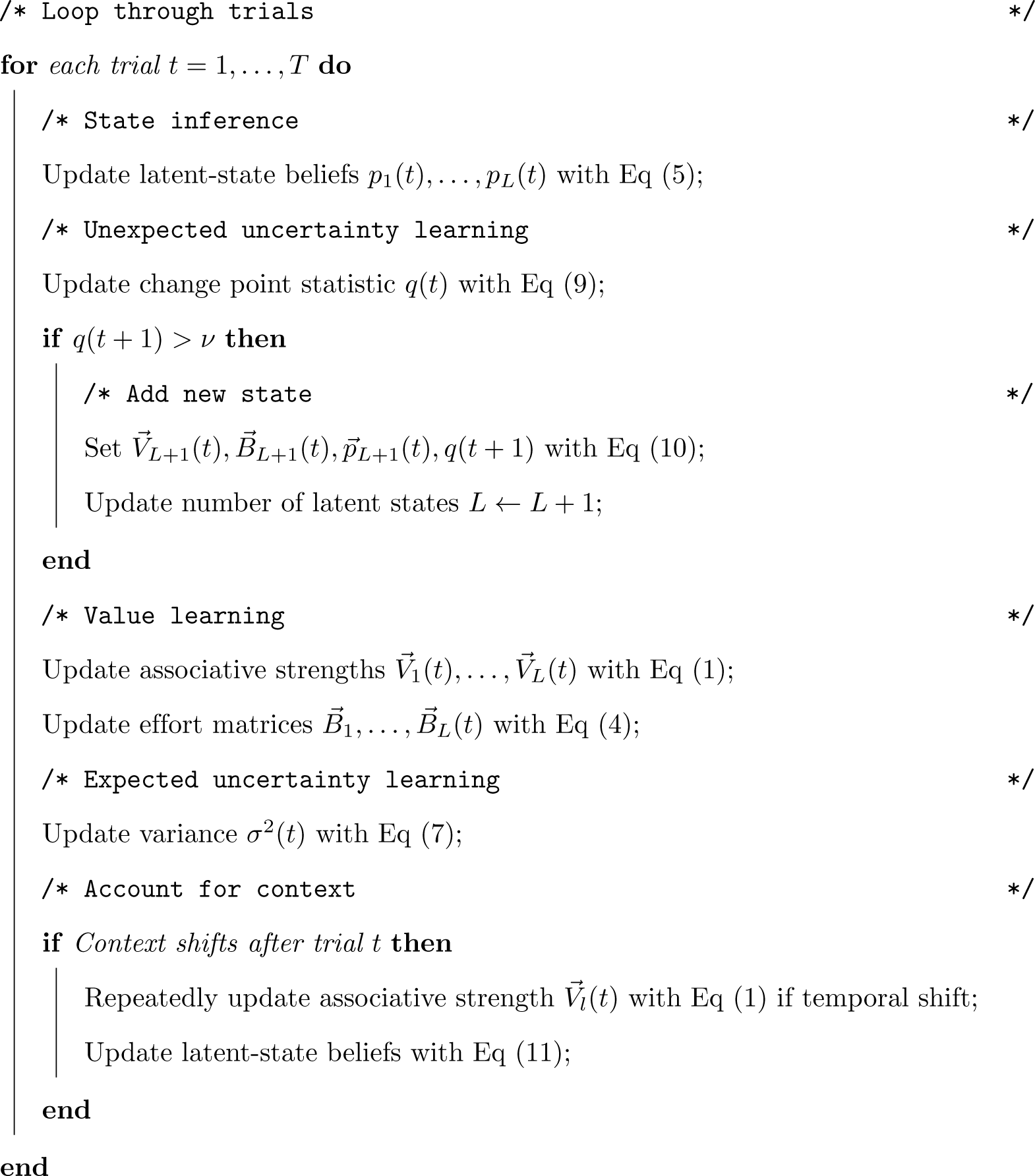

### Overview of simulation

We used simulation to verify that our latent-state model can reproduce a set of group-level effects observed from classical experiments on learning (Table 1). Our model was compared to the Rescorla-Wagner (RW) model, fRL+decay model of Niv et al [34] and three latent-state learning models: the Redish (2007) model [11], the infinite-mixture model of Gershman and Niv in [17], and the Gershman (2017) model in [19]. Other model comparisons are considered in S1 Text and S1-2 Figures. For each model, we computed associative strength per cue and belief in each latent state. For our model, the infinite-mixture model, and the Gershman (2017) model, we defined associative strength as the expected reward conditional on the cue being presented alone and belief in a latent state as the estimated probability of the latent state given observations. In order to compare the Redish (2007) model to other models, we defined associative strength for each cue as average estimated value between reinforcing and not reinforcing the cue presented alone and defined belief in a latent state to be 1 if the state was identified as the current agent state and 0 otherwise. Our model includes pairwise interactions between cues as additional cues and also centers rewards (i.e. *R*(*t*) = *−*1*/*2 and *R*(*t*) = 1*/*2 rather than *R*(*t*) = 0 and *R*(*t*) = 1). Additional details about the simulation are found in the Methods section.

**Table 1.** Simulated experiments and group-level effect observed in each experiment. Models were evaluated based on its ability to reproduce the observed effect for each simulated experiment.

| Experiment | Observed effect |
| --- | --- |
| Blocking [35] | Associative strength of a neutral cue is blocked when reinforced with a cue already associated with the reward |
| Overexpectation [36] | Associative strength of two cues weakens when these cues are reinforced together after reinforcing each cue separately |
| Conditioned inhibition [37] | A cue can inhibit responding to a reward after alternating between reinforcing a separate cue and presenting both cues with a weaker reward or no reward. |
| Wilson *et al* (1992) [38] | Associative strength of a cue increases more when the cue is a less accurate predictor of a reward (also known as the Hall-Pearce effect [24]) |
| Rescorla (2000) [39] | Changes in associative strength can be greater for one cue over another when presented at the same time |
| Partial reinforcement [40–42] | Associative strength of a cue is slower to extinguish after partial reinforcement compared to continuous reinforcement (also known as the partial reinforcement extinction effect) |
| Backwards blocking [43] | Associative strength of a cue weakens when a separate cue is reinforced after reinforcing both cues together |
| Rapid return of fear & renewal [12] | Associative strength of cue returns quickly after extinction, particularly when extinction is associated with a different context |
| Spontaneous recovery [12] | Associative strength of a cue returns more greatly after extinction with a longer interval between extinction and renewal |
| Memory modification [44, 45] | Associative strength of a cue to a fearful outcome can be weakened if extinction occurs within a certain time window following a single retrieval trial |

### Blocking and other effects

We first tested whether our model could reproduce learning effects that established the RW model as a powerful model of associative learning (Fig 2). In a blocking experiment, for instance, a learning agent is conditioned to associate a cue (Cue A) with a reward, leading to a high associative strength for Cue A. Afterwards, Cue A is repeatedly presented with another cue (Cue B). The compound is then reinforced with a reward. Even though Cue B is reinforced, Cue B does not acquire the high associative strength as Cue A did. That is, conditioning of Cue A *blocked* conditioning of cue B. The RW model, our latent-state model, and the Gershman (2017) model predict blocking because of the way associative strength is updated in each model: updates depend on prediction error and predictions depend on the sum of associative strengths of present cues. Consequently, associative strength of Cue B changes slightly, because the compound of an excitatory Cue A and neutral Cue B yields small prediction errors.

**Fig 2.**
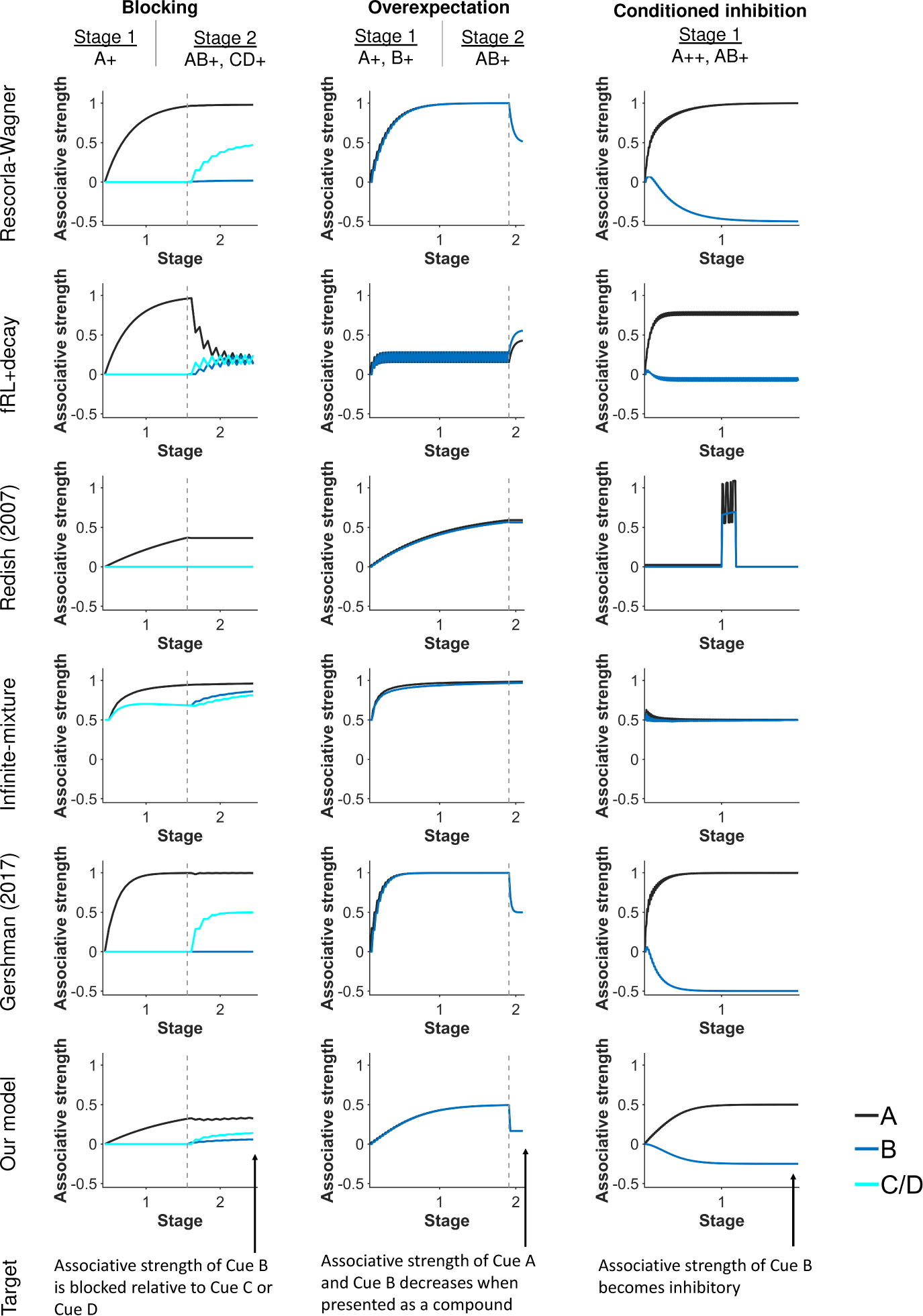
Simulated associative strengths of cues during blocking, overexpectation, and conditioned inhibition experiments, which the Rescorla-Wagner (RW) model famously could explain. Built upon the RW model, our latent-state model and the Gershman (2017) model can also explain these experiments. Gray dashed lines demarcate experimental stages.

Our latent-state model reproduces overexpectation and conditioned inhibition, two other group-level effects explained by the RW model. An overexpectation experiment consists of reinforcing two cues separately with a reward and then later as a compound. When presented as a compound, associative strengths decrease. A conditioned inhibition experiment consists of intermixing the reinforcement of one cue (Cue A) with the omission of a reward when the cue is presented with another cue (Cue B). Cue A increases in associative strength while Cue B decreases in associative strength. As with blocking, the RW model, our latent-state model, and the Gershman (2017) model capture overexpectation and conditioned inhibition because of the way associative strength is updated. For overexpectation, expectations are twice actual rewards when the cues are presented together, yielding negative prediction errors and a decrease in associative strength for each cue. For conditioned inhibition, the negative associative strength of Cue B negates the positive associative strength of Cue A so that the sum of their associative strengths predicts the weaker reward. The Redish (2007) and infinite-mixture models do not update associative strength in the same way as the RW model, whereas the fRL+decay model decays associative strength of cues that are not presented, leading to different predictions for blocking, overexpectation, or conditioned inhibition experiments.

### Learning effects not explained by the RW model

We also tested whether our latent-state learning model could reproduce historically-important learning effects not predicted by the RW model due to its assumption of constant associability (Fig 2).

#### Associability depends on beliefs in latent-states

Latent-state learning allows an agent to attribute sudden shifts in associations to new latent states, even when the shift is unsignalled. Associability starts relatively high when learning about a new state, thereby explaining the well-known Pearce-Hall effect [9] in which associability remains high in unpredictable circumstances.

To illustrate, we find that a shift in latent state beliefs can explain Experiment 1 from Wilson et al (1992) [38]. This experiment involved two groups, Group C and Group E. For Group C, the experiment consisted of alternating between reinforcing and not reinforcing two cues (a tone denoted by Cue B and a light denoted by Cue A) with a reward (food), followed by reinforcing just the light. Group E had similar experiment conditions, except the tone was omitted in the middle of the experiment on non-reinforced trials. The omission was expected to decrease the associative strength of the light for Group E relative to Group C, which would be observed when the light was paired with food. Surprisingly, Group E responded more favorably to the light than Group C. The leading explanation was that associability of the light was higher for Group E than Group C [10].

Our latent-state model offers an explanation: that while indeed the associative strength of the light does decrease for Group E relative to Group C, the pairing of the light (Cue A) solely with food is so surprising to Group E that they correctly infer the change in experimental conditions thereby shifting to a new latent state to build new expectations for the light (Fig 3). This shift would be observed as higher associability of the light in Group E than Group C, agreeing with experimental results. The Redish (2007) model also captures greater responding to the light in Group E than Group C. By contrast, the RW model, infinite-mixture, and Gershman (2017) models do not show greater responding to the light (cue A) in Group E. Meanwhile, the fRL+decay model captures this experiment but provides an alternative explanation: associative strength of the tone decays less in Group E because it is presented more often, which is compensated by an increase in the associative strength of the light.

**Fig 3.**
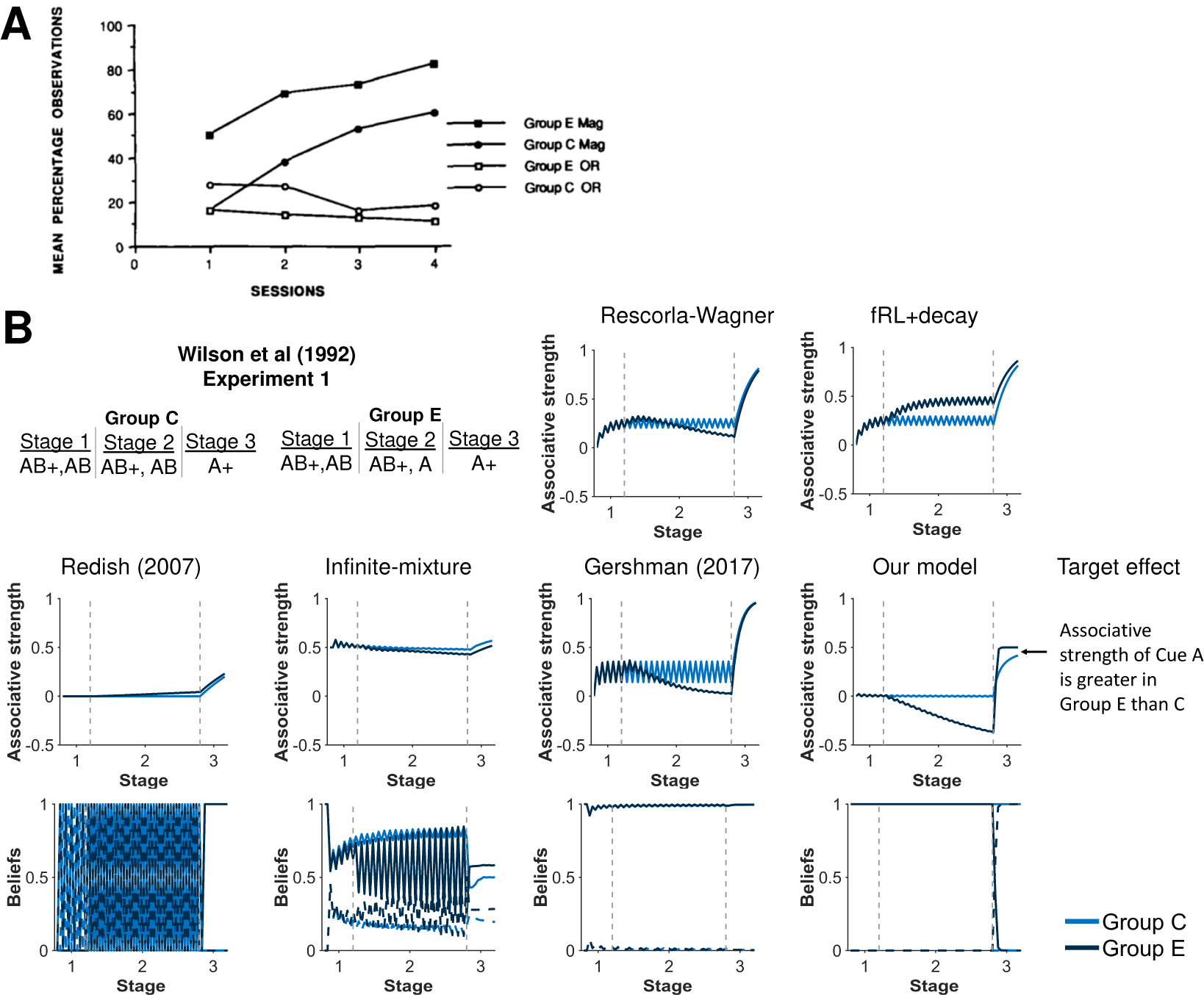
Associability depends on latent-state beliefs. **A)** Experimental results from Stage 3 of the Wilson et al (1992) experiment [27]. Group E had higher magazine activity, i.e. greater responding, than Group C during the light (Cue A) in Stage 3. Thus, it was believed the light had greater associative strength in Group E than Group C during Stage 3. Reprinted from Paul N. Wilson, Patrick Boumphrey, & John M. Pearce, Quarterly Journal of Experimental Psychology 44:1 pp. 17-36. Reprinted by Permission of SAGE Publications, Ltd. **B)** Simulation of associative strength of the light (Cue A) and latent state beliefs from the Wilson et al (1992) experiment. Our model predicts that only Group E detects the change in experimental conditions and shifts their beliefs. Because of this shift, associability is higher in Group E than C during Stage 3, leading to higher associative strength of the light. Beliefs in the first latent state (dark and light blue solid lines) and the second latent state (dark and light blue dashed lines) are shown for models with latent states. Gray dashed lines demarcate experimental stages.

Other learning effects can also be similarly explained by our model as a shift in latent-state beliefs (Fig 4). Experiments 1A-B in [39], for example, examined whether a cue (Cue A) would undergo the same change in associative strength as another cue (Cue B) when presented together, as would be predicted by the RW model. Rescorla, however, found that Cue A decreased more in associative strength than Cue B. Our latent-state model explains that differences in associability arises because of a shift in latent states. In Experiment 1B, for example, the omission of the reward is so surprising to the agent—Cue A was always reinforced—that learning agents correctly infer a change in conditions and switch their beliefs to a new latent state in which they do not expect Cues A and B to be reinforced. Since rewards were originally expected to follow Cue A but not cue B, the associative strength of Cue A experiences the greater change. By contrast, the only other models to capture greater associability of Cue B during stage 2 was the infinite-mixture model in experiment 1A and the Redish (2007) model in experiment 1B. Otherwise, the other models for the given set of parameters do not detect the change in experimental conditions, causing cues A and B to change a similar amount during Stage 2.

**Fig 4.**
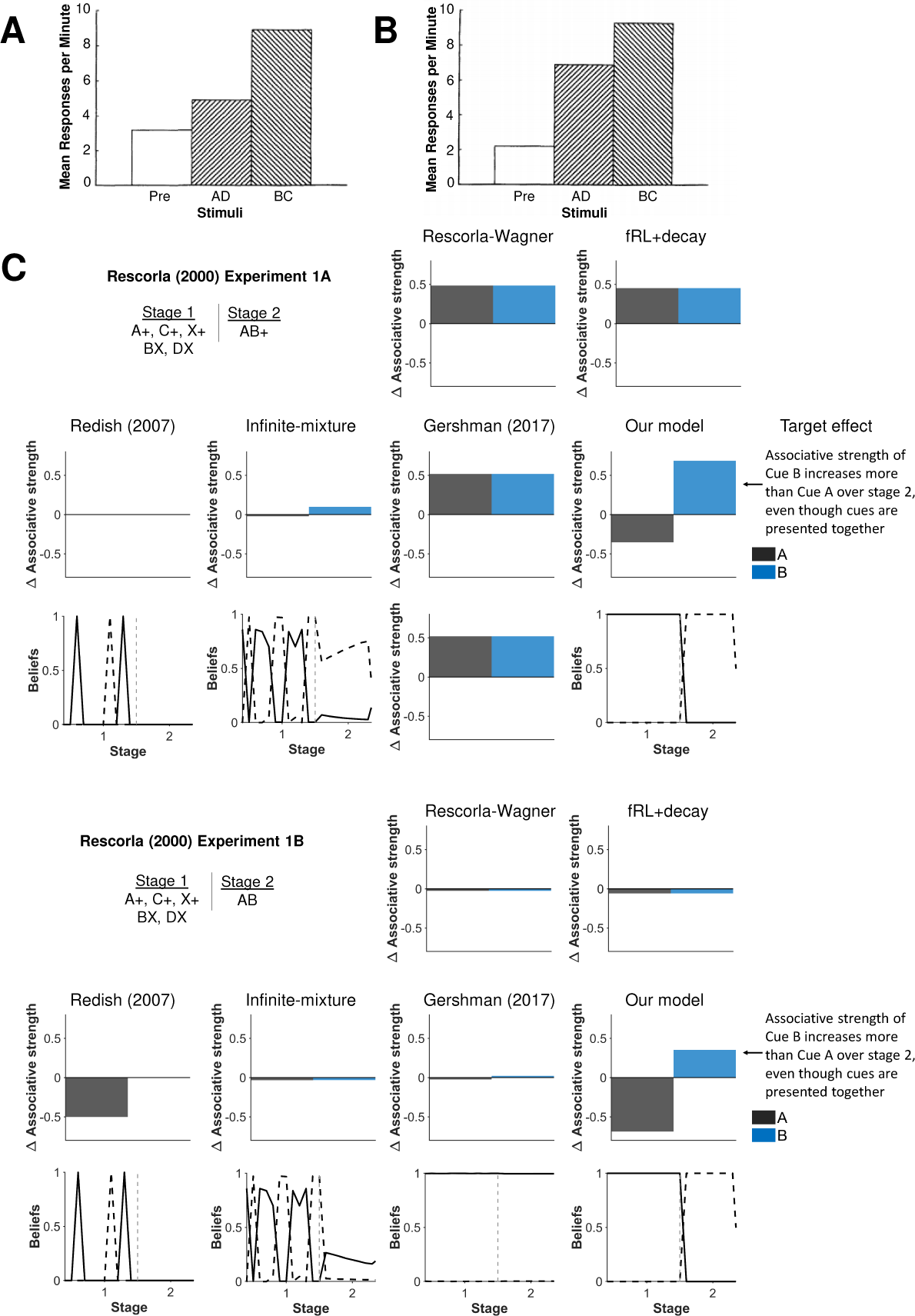
Another demonstration of associability depending on beliefs in latent states. Experimental results from **A)** Experiment 1A and **B)** Experiment 1B by Rescorla (2000) [39]. In both experiments, Rescorla concluded that associative strength increased more for Cue B relative to Cue A when presented together based on compound tests that showed greater responding in the compound with the Cue B than the compound with Cue A. This result suggested that associability can differ between cues even when presented together. Reprinted from “Associative Changes in Excitors and Inhibitors Differ When They Are Conditioned in Compound” by R.A. Rescorla, 2000, *Journal of Experimental Psychology*, *26*, p. 430-431. Reprinted with permission from the American Psychological Association. **C)** Total change in associative strength was simulated during Stage 2 of Experiments 1A-B from Rescorla (2000) [39]. The RW model does not capture these effects since associability is constant whereas our model captures these effects because latent-state beliefs alters associability. Beliefs in the first latent state (black solid lines) and the second latent state (black dashed lines) are shown for models with latent states. Gray dashed lines demarcate experimental stages.

The partial extinction learning effect (PREE) is also similarly explained by our model as a shift in latent-state beliefs (Fig 5). Partial reinforcement involves alternating between reinforcing and not reinforcing a cue. It was found that a response was harder to extinguish after partial reinforcement compared to continuous reinforcement [40]. Our model explains that an agent is better able to discriminate between reinforcement and extinction with a continuous reinforcement schedule. This allows the agent to switch their beliefs to a new latent state and thus experience faster extinction. PREE was also observed even if continuous reinforcement was administered in between partial reinforcement and extinction [41, 42]. In this case, an agent is still better able to discriminate between reinforcement and extinction with a continuous reinforcement schedule, but differences between reinforcement schedules are smaller as an agent shifts their beliefs to a new latent state in both cases. Only the Redish (2007) model was also able to capture greater extinction for continuous reinforcement compared to partial reinforcement in either of the two PREE simulation experiments.

**Fig 5.**
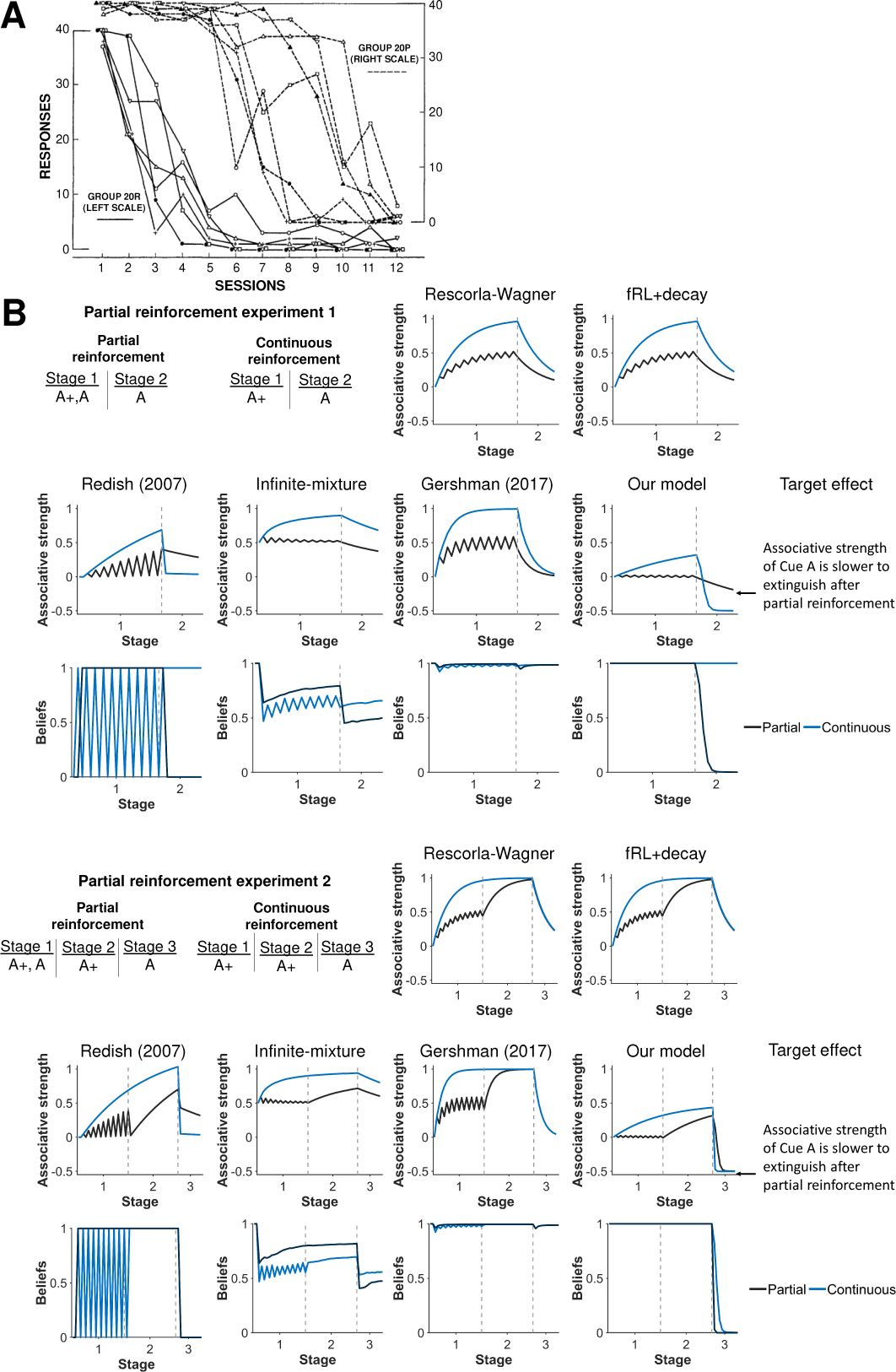
Partial reinforcement extinction effect. **A)** Experimental results from Jenkins [41] demonstrating that the associative strength of a cue is harder to extinguish after partial reinforcement (Group 20P) compared to continuous reinforcement (Group 20R). Reprinted from “Resistance to extinction when partial reinforcement is followed by regular reinforcement” by H.M. Jenkins, 1962, *Journal of Experimental Psychology*, *64*, p. 443. Reprinted with permission from the American Psychological Association. **B**) Simulation of partial reinforcement effect (Experiment 1). This effect is observed even when partial reinforcement is followed by continuous reinforcement prior to extinction (Experiment 2). Our model captures these effects, because an agent is better able to discriminate between reinforcement and extinction with a continuous reinforcement schedule. The agent can thus shift their beliefs to a new latent state in order to build new associations for extinction. Beliefs in the first latent state are shown for models with latent states. Gray dashed lines demarcate experimental stages.

#### Associability depends on history of cue presentation

Our model encodes the history of cue presentation in effort matrices *B_l_*(*t*), endowing associability in our model with unique properties. One example is backwards blocking (Fig 6). A compound of two cues (Cues A and X) is reinforced with a reward followed by only one of the cues (Cue X) being reinforced. In the second part, Cue A gains associative strength while Cue X loses associative strength *even though it is not presented*. The RW model cannot explain backwards blocking, because a cue must be present to change its associative strength. The infinite-mixture and the Gershman (2017) model also did not capture backwards blocking. By contrast, the associative strength of Cue X correctly decreased in the second part for our latent-state model. Our latent-state model first learns the compound predicts a reward. Without additional information, it splits associative strength equally between the two cues. Later, our latent-state model learns Cue A predicts rewards. Reconciling both parts, our latent-state model increases the associative strength of Cue A while decreasing the associative strength of Cue X. In other words, our latent-state model learns about the *difference* in associative strengths between Cues A and X. Mathematically, applying the inverse of *B_l_*(*t*) to the cue vector *c*(*t*) rotates the cue vector from being in the direction of Cue A to be in the direction of Cue A minus Cue X. Only the fRL+decay model was also able to reproduce backwards blocking, but provided an alternative explanation: associative strength decays for any cue that is absent, such as Cue X in Stage 2 of this experiment.

**Fig 6.**
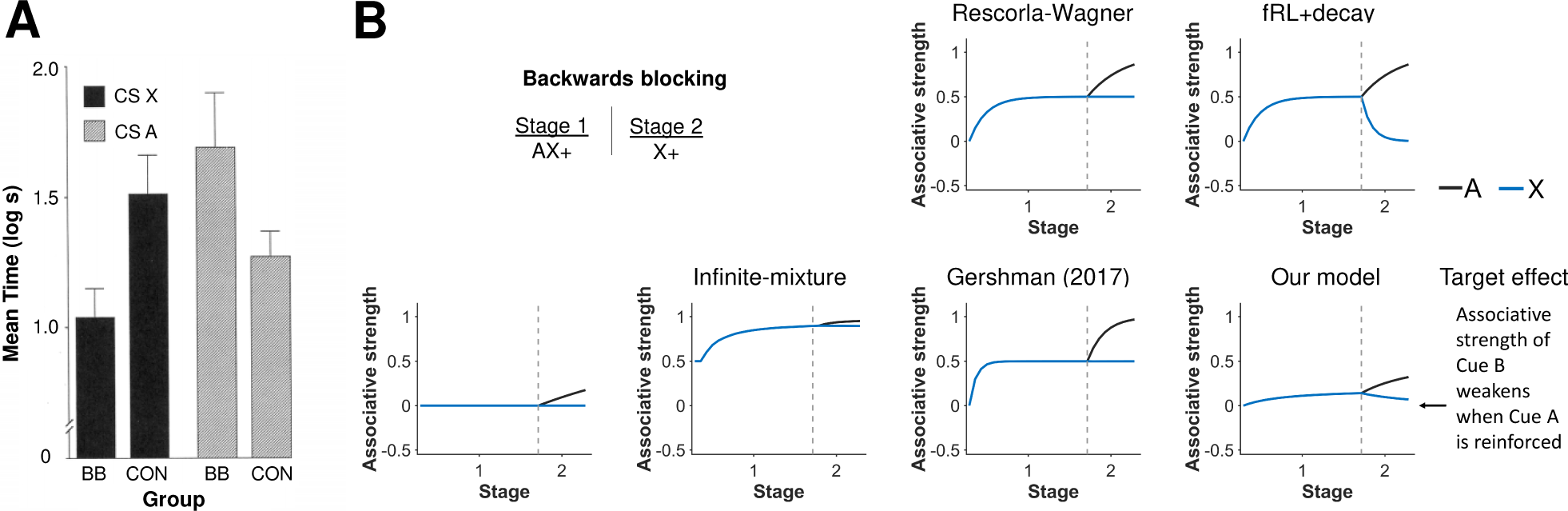
Associability depends on history of cue presentation. **A)** Experimental results of a backwards blocking experiment from Miller and Matute [43]. After two stages of the experiment, a backwards blocking group (BB) had significantly slower response time to Cue X than a control group (CON) even though Cue X was presented in an identical manner between groups. Their result suggested the associative strength of Cue X can change on trials it is not presented. Reprinted from “Biological Significance in Forward and Backward Blocking Discrepancy Between Animal Conditioning and Human Causal Judgement” by R.R. Miller and H. Matute, 1996, *Journal of Experimental Psychology*, *125*, p. 374. Reprinted with permission from the American Psychological Association. **B)** Simulation of a backwards blocking experiment. In our model, associability depends on the history of cue presentation through effort matrices. After the combined associative strength of Cue A and Cue X is learned, these effort matrices rotate the direction of learning into the direction of the difference of Cue A and Cue X. As a result, associative strength of Cue X decreases even though Cue A is presented alone, thereby allowing our model to capture backwards blocking. Gray dashed lines demarcate experimental stages.

### Learning with latent-states

To complete our simulation study, we tested if our latent-state model could describe more recent experiments that examine latent-state learning.

#### Renewal

Latent-state learning was offered as an explanation of renewal of expectations after extinction and the role that context plays in this renewal [11, 12, 15, 16]. We simulated renewal wherein a cue is reinforced, extinguished, and then reinforced (Fig 7). Two experimental conditions are considered: one in which the same visual/spatial context is provided through each phase and another in which a different visual/spatial context is provided during the extinction phase. Following prior models [8, 11, 17, 19], context was encoded as a separate cue in all models except our latent-state model. Our latent-state model encodes context as a shift in beliefs. Thus, while context may directly modulate both associative strengths and latent state inferences in other models, it only modulates latent state inferences in our model. This latter view is consistent with Bouton [12] who says “contexts modulate or ‘set the occasion’ for the current CS–US or CS–no US association. Put another way, they activate or retrieve the current relation of CS with the US.” Note for this example, expected rewards are given on each trial to account for the contribution of both the cue and the context rather than associative strength which only accounts for the contribution of the cue.

**Fig 7.**
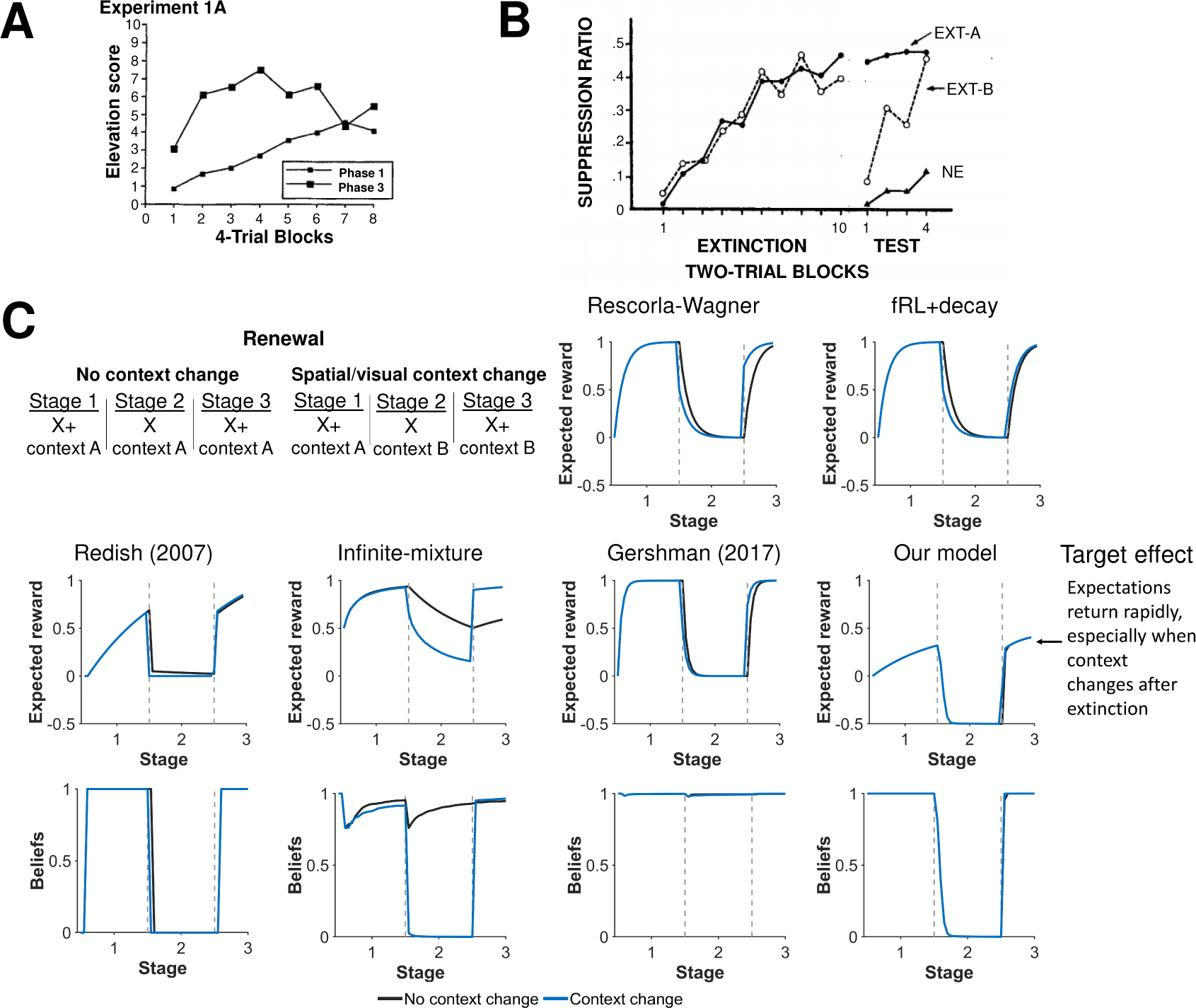
Changes in context influences beliefs. **A)** Experimental results from Ricker and Bouton [46] demonstrating a faster response during reacquistion (phase 3) than acquisition (phase 1) after extinction (phase 2). Reprinted by permission from Springer Nature Customer Service Centre GmbH: Springer Nature, Animal learning and behavior, Reacquisition following extinction in appetitive conditioning, Sean T. Ricker & Mark E. Bouton, (1996). **B)** Experimental results from Bouton and King [47] demonstrating a more robust return of a fear when extinction occurs in different context (EXT-B) as opposed to the same context (EXT-A) as acquisition. Reprinted from ‘Contextual Control of the Extinction of Conditioned Fear: Tests for the Associative Value of the Context” by M.E. Bouton and D.A. King, 1983, *Journal of Experimental Psychology: Animal Behavior Processes*, *9*, p. 252. Reprinted with permission from the American Psychological Association. **C)** Simulation results of expected rewards when a response is reinstated after extinction with and without a change in a visual/spatial context. The second experiment examines associative strength of cue when a response is reinstated after extinction with and without a change in a temporal context (i.e. a time delay between trials). Our model shows a rapid reinstatement of expectations as the agent switch their beliefs back to the first latent state. Our model also shows that rapid reinstatement is more robust with changes in context, particular in the first few trials of reinstatement. Expected rewards are depicted rather than associative strengths to account for the influence of context on expectations in addition to the cue, since models other than our model treat context as an additional cue. Beliefs are shown for the first latent state. Gray dashed lines demarcate experimental stages.

In both experimental conditions, our latent-state learner slowly detects the shift when arm-reward contingencies first shift and switches their beliefs to a new latent state, effectively consolidating the memory of the first task phase. By detecting this shift, the agent can both construct new associative strengths for each arm and preserve the old associative strengths that were accurate for the first block of trials. These associative strengths are later *recalled* when arm-reward contingencies revert back and the agent switches their beliefs back to the original latent state. Introducing a different context during extinction allowed the learning agent to better detect the underlying structure of the task, recalling the first latent state earlier during reinstatement, resulting in higher associative strength in the final phase. This improvement in detection predicted by the model, however, was very slight and most noticeable in the first few renewal trials. The infinite-mixture model predicts that the learning agent switches beliefs in a new latent state only when a different context is introduced otherwise using one latent state to predict all phases of the task. For the given parameters, the Gershman (2017) model uses only one latent state for all three phases of the task for both the no renewal and renewal conditions. Consequently, other than our model, only the Redish (2007) model correctly predicts a rapid return of expectations.

#### Spontaneous recovery

Latent-state learning also provides an explanation for how changes in temporal context influences reinstatement (Fig 8). For example, spontaneous recovery is a learning effect wherein the associative strength of a cue is reinstated more strongly after a time delay between extinction and renewal [12]. We simulated spontaneous recovery using the same schedule as renewal, but adding a time delay between extinction and renewal. Our model accounts for time delays by shifting beliefs towards uniform beliefs over latent states. Uniform beliefs reflects that beliefs from a prior trial is less informative on a trial when there is a time delay between trials. As a result, a time delay causes the associative strength of the cue predicted by our model to be about half the associative strength at the end of its initial acquisition (first latent state) and at the end of extinction (second latent state). Without the time delay, the associative strength predicted by our model is simply the associative strength at the end of extinction. Only the Gershman (2017) model adjusts its prediction for context changes due to temporal shifts, but does not capture a more robust return of associative strength due to a temporal shift. The other models also do not capture this more robust return.

**Fig 8.**
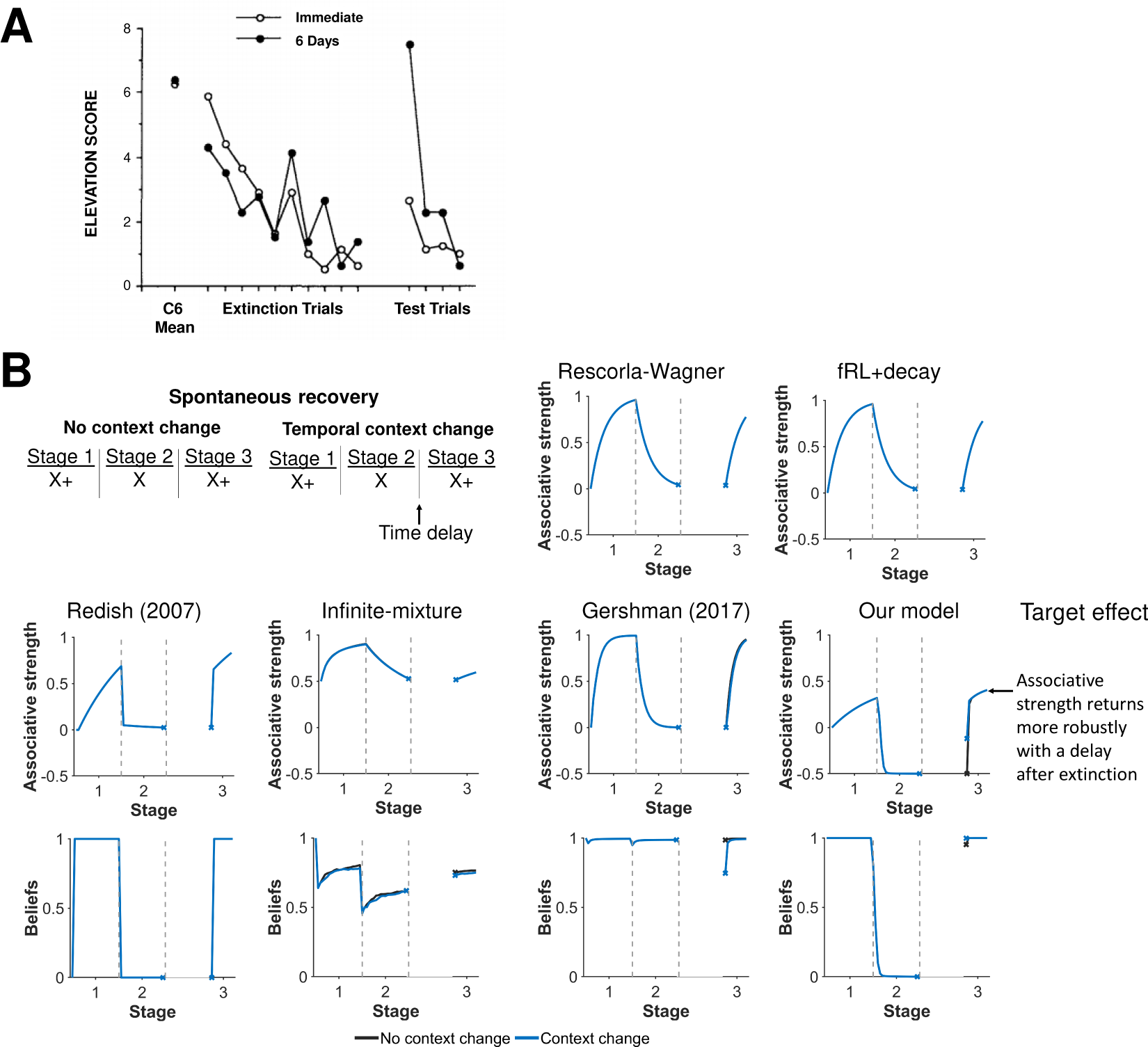
Changes in temporal context also influences beliefs. **A)** Experimental results from Brooks and King [48] demonstrating a more robust return of a fear when extinction occurs after a 6 day delay as opposed to immediately after acquisition. Reprinted from “A Retrieval Cue for Extinction Attenuates Spontaneous Recovery” by D.C. Brooks and M.E. Bouton, 1993, *Journal of Experimental Psychology: Animal Behavior Processes*, *19*, p. 80. Reprinted with permission from the American Psychological Association. **B)** Simulation results of associative strength and beliefs when a response is reinstated after extinction with and without a change in a temporal context (i.e. a time delay between trials). Our model shows rapid reinstatement is more robust with changes in temporal context, particular in the first few trials of reinstatement. Beliefs are shown for the first latent state. Gray dashed lines demarcate experimental stages.

#### Memory modification

Our last simulation experiment explores the potential role of latent-state learning in memory modification. Monfils et al [44] in a rodent model and later Schiller et al [45] in humans showed that a single *retrieval* trial between acquisition and extinction can reduce the fear response to the cue in latter tests. The time between retrieval and extinction was further shown to modify this effect. Only certain windows of times reduced the fear response. They proposed that the retrieval trial helped to reconsolidate the acquisition memory to be less fearful within this *reconsolidation* window. The Gershman (2017) model [19] includes a computational explanation of this phenomenon. With a similar Bayesian framework to the Gershman (2017) model, our model included similar memory modifications.

We simulated the Monfils-Schiller experiment in [19], varying the time between retrieval and extinction (Fig 9). Our model reproduces a qualitatively similar result as the Gershman (2017) model. There is a reconsolidation window which can help reduce the fear memory. For example in both models, associative strength of the cue during testing is smaller for a time of 5 from retrieval to extinction relative to time of 1 or 100. Both models predict that beliefs in the first latent state (corresponding primarily with acquisition) remains relative high in the reconsolidation window, allowing the agent to update the acquisition memory to be less fearful. Outside the reconsolidation window, the agent updates the second latent state (corresponding primarily with extinction), leaving the acquisition memory relatively intact. While qualitatively similar between models, the effect of retrieval is more pronounced in our model in terms of differences in magnitude of cue associative strength during testing.

**Fig 9.**
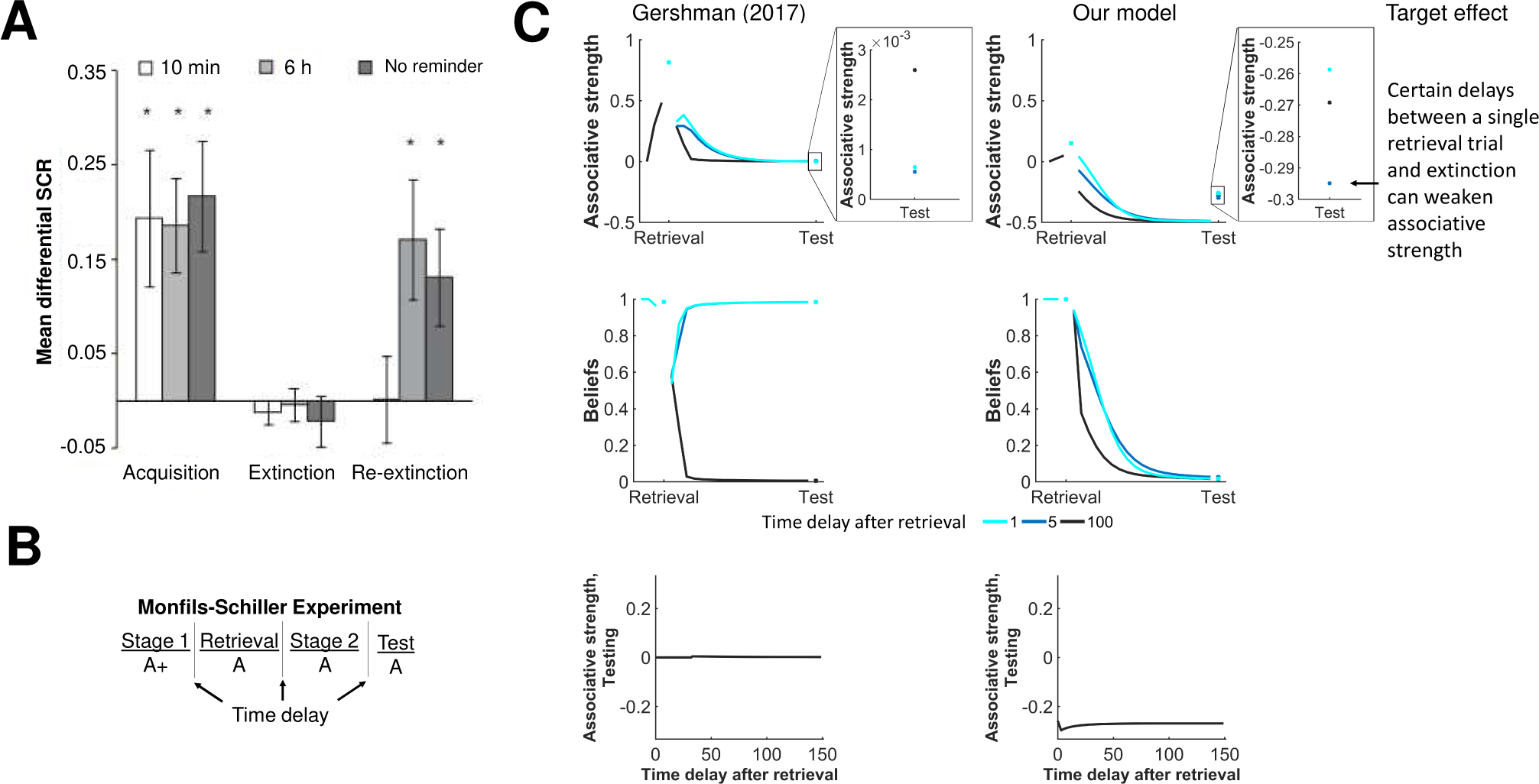
Memory modification. **A)** Experimental results from Schiller et al [45] demonstrating that different delays (10 min, 6 hr, no reminder) of a retrieval trial can significantly modify fear response after extinction. Reprinted by permission from Springer Nature Customer Service Centre GmbH: Springer Nature, Nature Preventing the return of fear in humans using reconsolidation update mechanisms, Schiller et al., (2010). **B-C)** Simulation results of associative strength and beliefs for three different time delays (1,5, and 100) between a single retrieval and extinction. Both models correctly predict that certain time delays can weaken the associative strength upon testing. The bottom row depicts a ‘reconsolidation window’ of time delays after retrieval wherein the associative strength on testing is decreased.

### Overview of simulation results

Our model reproduced group-level effects from a series of classical experiments. Table 2 summarizes which effects were captured by which model. Different aspects of the model are important for reproducing different effects. Updating associative strength similar to the RW model, for example, allows our model to capture effects famously reproduced by the RW model (e.g., blocking). Updating associability based on latent-state beliefs captures sharp increases in associability when a learner shifts their beliefs to a latent state, reproducing experiments from Wilson et al (1992) [38] and Rescorla (2000) [39], and experiments involving partial reinforcement [40–42]. Additionally, updating associability based on history of cue presentation rotates updates in different directions, allowing our model to reproduce backwards blocking. Further, using latent-states to index disparate cue-reward associations allows for a rapid return of prior expectations as a learner recalls a prior latent-state. Changing latent-state beliefs to reflect contextual changes allows our model to capture effects modulated by visual or temporal context such as renewal or spontaneous recovery, and together with rumination steps, allows our model to reproduce memory consolidation experiments [44, 45]. Finally, our model uses interaction terms and centers rewards, which we found were important for explaining certain group-level effects by determining whether a learning agent would shift to a new latent state.

**Table 2.** Models were assessed based on whether or not they could reproduce the observed effect for each simulated experiment (yes=✓ or no=✗), with observed effects defined in Table 1. It is important to note, however, that this assessment does not account for the magnitude of the effect or whether a different set of parameters could reproduce the effect.

| Experiment | RW | fRL+decay | Redish | Infinite-mixture | Gershman | Our model |
| --- | --- | --- | --- | --- | --- | --- |
| Blocking | ✓ | ✗ | ✗ | ✗ | ✓ | ✓ |
| Overexpectation | ✓ | ✗ | ✗ | ✗ | ✓ | ✓ |
| Conditioned inhibition | ✓ | ✓ | ✗ | ✗ | ✓ | ✓ |
| Wilson *et al* (1992) | ✗ | ✓ | ✓ | ✗ | ✗ | ✓ |
| Rescorla (2000) Exp 1A | ✗ | ✗ | ✗ | ✓ | ✗ | ✓ |
| Rescorla (2000) Exp 1B | ✗ | ✗ | ✓ | ✗ | ✗ | ✓ |
| Partial reinforcement | ✗ | ✗ | ✓ | ✗ | ✗ | ✓ |
| Backwards blocking | ✗ | ✓ | ✗ | ✗ | ✗ | ✓ |
| Rapid return of fear & renewal | ✗ | ✗ | ✓ | ✗ | ✗ | ✓ |
| Spontaneous recovery | ✗ | ✗ | ✗ | ✗ | ✗ | ✓ |
| Memory modification | ✗ | ✗ | ✗ | ✗ | ✓ | ✓ |

## Discussion

We presented a computational model of latent-state learning, using the Rescorla-Wagner (RW) model as its foundation, and positing that latent states represent disparate associations between cues and rewards. In a series of simulation experiments, we tested the model’s ability to reproduce learning experiments both famously explained and not explained by the RW model and to capture behavior in recent experiments testing latent-state inferences. Lastly, we formally derived our latent-state model under the premise of computational rationality, i.e. a learning agent wants to predict outcomes efficiently.

Our goal was to identify a model of latent-state learning that reproduces group-level effects from classical conditioning experiments. The resulting model makes five critical assumptions/predictions about how an agent learns cue-reward associations: an agent learns online, an agent uses latent states to predict rewards, associability depends on beliefs in latent states and cue novelty, latent states are relatively stable between trials, and beliefs are maintained over multiple latent states. Including these features helps to ensure that the proposed model could examine learning in numerous and diverse experiments, from experiments of classical conditioning to more recent experiments on latent-state inferences. We show that most features fall-out naturally when trying to develop an online approach to reward prediction which uses latent-state inferences. Other models of latent-state learning share some, but not all these features [11, 17, 19].

For example, our model assumes a learning agent uses latent states to index disparate *associations*, or mappings, from cues to rewards in an effort to predict cues. By contrast, the Redish (2007) and infinite-mixture models use latent states to index disparate *combined observations* of cues and rewards, which can in turn be used to infer rewards from a set of cues. Also, our model assumes latent states evolve according to a Markov chain, whereas other models assume latent states are exchangeable or independent between trials. As a consequence, beliefs in latent states are relatively stable between trials. This stability was observed in certain simulation experiments, in which the predominant latent state would switch only one or two times for our model but over a dozen times for the infinite-mixture model and Redish (2007) model.

Interestingly, our model still updates associative strength in a similar manner to a RW model: change in associative strength is prediction error scaled by associability. In fact, the RW model is a specific case of our latent-state model when there is only one latent state and associability is fixed. For this reason, our latent-state model can reproduce similar learning effects as the RW model such as blocking, overexpectation, and conditioned inhibition. Some latent-state models cannot capture all of these classic learning features [11, 17]. However, our latent-state model goes beyond the RW model by allowing associability to vary across trials. This feature is thought to be important for describing learning effects not explained by the RW model, such as the Pearce-Hall effect, and is incorporated into newer models, such as the Hybrid RW model [9, 10, 49]. Associability in our latent-state model involves two components: effort matrices and beliefs in latent-states. Because associability depends on effort matrices, changes in associative strength are rotated into a direction which has not been previously learned, helping to explain backwards blocking. These matrices approximate Fisher information matrices, which are often incorporated into iterative approaches to maximum likelihood estimation [28, 50]. Meanwhile, because associability also depends on beliefs, associative strength can be updated quickly when a learning agent infers a sudden shift in experimental conditions. This feature allowed our latent-state model to capture experimental results from Experiment 1 in Wilson et al (1992) [27] and Experiments 1A–B in Rescorla (2000) [39].

Similar to other latent-state models, our model also assumes an agent who learns online. Online learning requires only current observations rather than the entire history of observations to update parameters and quantities such as associative strength. Consequently, online approaches have computational and memory requirements that grow in the number of latent states rather than in trials, which could be a more realistic reflection of how humans learn. The infinite-mixture model also uses online updates for all its variables [17], whereas the Gershman (2017) model and Redish model (2007) uses online updates for most variables [11, 19]. For comparison, an optimal Bayesian approach to latent-state learning would require enumerating over all possible sequences of latent-states, a number that grows exponentially with each trial, imposing a computational and memory burden that is believed to be overly optimistic for human computation [26].

Critically, our latent-state model relies on an approximate Bayesian filter to maintain and update beliefs over latent-states online. Other models of latent-state learning use other approaches to reduce computational burden such serial representations of committed beliefs rather than distributing belief across multiple hypotheses [11], particle methods [15, 17, 19], or maximum a posteriori estimates [19]. Additional assumptions might also be used such as exchangeable trials and binary outcomes [17], one latent state [34], one latent-state transition [27], or known cue-reward associations [32]. Assumptions, however, limit the ability of any model to describe learning in varied experiments. For example, we can capture renewal, spontaneous recovery, and other learning phenomena that require a more general number of latent states and transitions because of our use of an approximate Bayesian filter [28]. We can also capture experiments such as [32] that require beliefs to be maintained over over multiple latent-states. Notably, the experiment [32] suggested a link between log-beliefs and activity in the orbitofrontal cortex.

Many of the assumptions/predictions of the presented model are testable. For example, associability for a latent-state is predicted to be directly dependent on the beliefs in that latent-state. Stronger beliefs lead to faster learning, whereas weaker beliefs lead to slower learning. An experiment could be designed to test, for example, if associability decreases by 50% when beliefs are split between two latent-states relative to when beliefs were concentrated on one latent-state. Further, associability is predicted to depend on whether cues had previously been presented together. New combination of cues can lead to quick learning even when individual cues making up the combination have already been presented. Last, latent-state beliefs are predicted to be updated in a non-optimal manner, suggesting one try to induce sub-optimal behavior in an experiment.

Several limitations of the proposed model should be considered. First, we anticipate there are group-level learning effects that our model fails to describe or there are other models that could explain participant behavior. Second, we did not examine to what extent beliefs in latent-states and other variables from our model are encoded in various brain regions. Third, we assume beliefs are maintained over multiple latent states, but some experiments suggest that an agent is committed to a state and alternates between committed states [33]. One possibility is that beliefs in latent states influence which state is committed and over time, the frequency of committed states mirrors these beliefs. Modalities such as fMRI might then suggest beliefs are maintained over multiple states, as in [32], because they lack the temporal resolution needed to detect the alternating committed states. Allowing beliefs to determine committed states might even lead to better explanations of participant data, as it allows for more flexibility in how states are updated for a given participant. Fourth, there may be better ways to integrate computational mechanisms of how memories are formed into our model [51]. For example, we use online updates in order to avoid memory requirements that grow in the number of trials, but it is possible that humans are either capable of such requirements or use other ways than online updates to integrate past observations. Fifth, some effects (e.g., memory modification effect [44, 45]) reproduced by our model might be too subtle to ever be detected experimentally. Sixth, we assume latent states are relatively stable between trials. This assumption may be accurate for describing learning in experiments, since experiments often involve blocks of trials in which rewards/cues are generated by the same rules, but may be less accurate for real-world learning environments. Seventh, we included interactions between cues to influence when a learning agent shifts to a new latent state, which we found was necessary to recover one of the Rescorla (2000) experiments [39]. This inclusion, however, should be investigated further to determine if it reflects how agents actually learn. Finally, we did not study model performance in terms of predicting rewards as a consequence of using an approximate Bayesian filter. It may useful to know how far from optimal our approximate method is for maximum likelihood estimation in various settings.

In sum, this work establishes the validity of an online model for latent-state inferences that generalizes across experiments. Establishing such a framework is a necessary step towards quantifying normative and pathological ranges of latent-state inferences (and mediating neurocircuitry) across varied contexts. Overall, this work moves us closer to a precise and mechanistic understanding of how humans infer latent-states.

## Methods

### Simulation details

Parameters were fixed throughout the simulation (Table 3). Parameters for alternative models were chosen from their respective papers, except for the maximum number of latent states which we fixed at 15 to reduce memory requirements. Our model includes cues throughout simulation to represent pairwise interaction terms (i.e. a binary indicator if a pair of cues are present or absent). Our model also centers rewards with *R*(*t*) = 1*/*2 when a reward is presented and *R*(*t*) = *−*1*/*2 when a reward is not presented. That way, initial associative strengths give rise to neutral (i.e. 50-50) expectations for reward. Thus, associative strengths are shifted by 1/2 for our model when compared to other models. Accompanying code is publicly-available at https://github.com/cochran4/OnlineLatentStateLearning. S1 Text provides further details on the schedules for how cues and rewards were delivered during each simulation task and examines sensitivity of our model’s predictions to changes in parameters.

**Table 3.** Fixed set of parameters used for all simulated experiments.

| Model | Parameter | Description | Value |
| --- | --- | --- | --- |
| Our model | $\alpha_0$ | Associative strength learning rate | 0.05 |
| Our model | $\beta_0$ | Variance learning rate | 0.05 |
| Our model | $\gamma$ | Latent-state transitions | 0.05 |
| Our model | $\sigma_0$ | Initial expected uncertainty | 0.5 |
| Our model | $\nu$ | Threshold for new state | 0.2 |
| Our model | $\delta$ | Unexpected uncertainty update | 0.6 |
| Our model | $\chi$ | Rumination steps | 5 |
| Rescorla-Wagner | $\alpha$ | Learning rate | 0.15 |
| Infinite-mixture | $\alpha$ | Concentration parameter | 1 |
| Infinite-mixture | $M$ | Number of particles | 100 |
| Gershman (2017) | $\alpha$ | Concentration parameter | 0.1 |
| Gershman (2017) | $\eta$ | Learning rate | 0.3 |
| Gershman (2017) | $\sigma_r^2$ | Reward variance | 0.4 |
| Gershman (2017) | $\sigma_x^2$ | Cue variance | 1 |
| Gershman (2017) | n/a | Rumination steps | 3 |

### Mathematical justification

We justify our choice of model under the premise of *computational rationality* [52], i.e. a learning agent wants to predict rewards efficiently. An optimistic strategy for the learning agent would be to use maximize likelihood estimation (MLE) to aid prediction. This would entail starting with a probabilistic model of rewards defined up to an unknown parameter *θ* and finding *θ* to maximize the log-likelihood of rewards (scaled by 1*/t*):

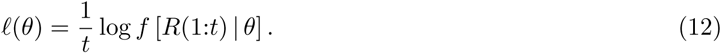

Treating rewards as a continuous variable, we use *f* to denote a probability density function. The estimate of *θ* together with the reward model predict future rewards. With an eye towards efficiency, we focus on *online* approaches, such as the Rescorla-Wagner model, and refer the reader to the work in [50] and [28] on updating latent-variable models online.

#### Rescorla-Wagner model as an online approach to maximum likelihood estimation

Before we motivate our latent-state model, we explore how one could perform MLE online to predict rewards without latent-states. In the process, we show that the RW model is an online approach to MLE and illuminate a pathway for performing MLE online with latent-states. The most natural modeling choice for rewards is arguably a standard Gaussian regression model, ubiquitous in statistical inference:

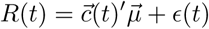

where 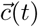 is the cue vector on trial *t* (independent variables); 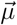 are unknown parameters (unknown regression coefficients); and *ε*(*t*) is an independent Gaussian random variable (error) with mean zero and unknown variance *ν*. Unknown parameters are collected in 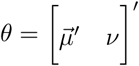.

Although we could maximize ℓ directly, we want an online approach to maximization and instead consider applying Newton’s method for optimization (i.e., gradient descent), which involves repeatedly updating an estimate 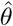 of *θ* with a new estimate of the form:

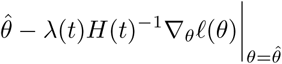

for some learning rate *λ*(*t*) and an appropriate matrix *H*(*t*). Matrix *H*(*t*) can take many forms and is important for determining how quickly estimates converge to a maximum/minimum. An ideal choice is the Hessian resulting in the Newton-Raphson method, but the Hessian is often noninvertible in MLE. A common alternative is the negative of the Fisher information matrix *I*(*θ*) scaled by 1*/t* and evaluated at the current estimate [50]. Even with Newton’s method, rewards still need to be stored up to trial *t*, but Newton’s method can be turned into an online algorithm using the well-known Robbins-Monro method [53]. We simply replace the gradient with an (stochastic) approximation that depends only on the current trial:

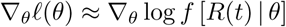

Note that each side of the equation is equal in expectation when rewards are independent and identically-distributed.

Upon replacing the gradient with a stochastic approximation, we arrive at the RW model if we use a constant learning rate *λ*(*t*) = *λ*_0_ and the matrix

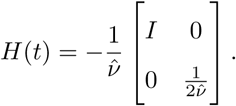

For the Gaussian regression model, matrix *H*(*t*) resembles

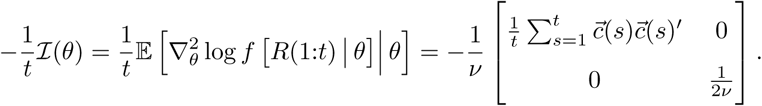

evaluated at 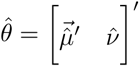 We simply replaced 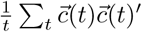 with the identity matrix *I*, which ensures *H*(*t*) is invertible. Putting the online approach together with the fact that

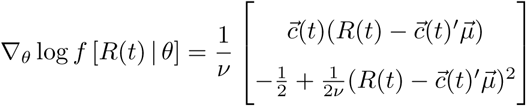

for the Gaussian regression model, we arrive at the following online update for the estimate 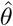:

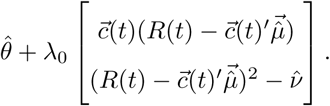

Critically, estimates 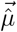 are updated exactly as in an RW model:

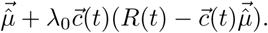

In other words, the RW model is an online approach to maximum likelihood estimation.

#### Latent-state learning as an online approach to expectation-maximization

Motivated by the previous argument, we now explore how one could perform MLE online to predict rewards with latent-states. A natural choice for modeling rewards is to use a Markov chain of Gaussian regression models. Latent random variable *X*(*t*) *∈ {*1*, …, L}* represents the active hypothesis on trial *t*. Given the active hypothesis *X*(*t*) = *l*, rewards are described by a Gaussian linear regression model:

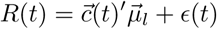

Where 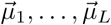 are unknown parameters and *ε*(*t*) is an independent Gaussian random variable with mean zero and unknown variance *ν*. The extension to latent-state learning thus leads to *L* different Gaussian regression models for rewards rather than one. As discussed in the Model subsection, we assume *X*(*t*) is a Markov chain that transitions to itself from one trial to the next with probability 1 *− γ*(*L −* 1)*/L* and transitions to a new state (all new states being equally-likely) with probability *γ*(*L −* 1)*/L*. Parameter *γ* and initial probabilities for *X*(0) are assumed to be known/given, leaving unknown parameters collected in 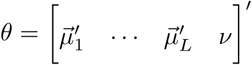.

In a latent-variable setting, a popular algorithm to perform MLE is expectation-maximization (EM) [54]. An iterative algorithm, EM improves upon a current estimate 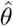 at each iteration in two steps. Fixing the current trial number *t*, an *expectation* step uses a current estimate 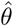 to calculate the expectation of the complete log-likelihood of rewards and latent states given rewards and scaled by 1*/t*:

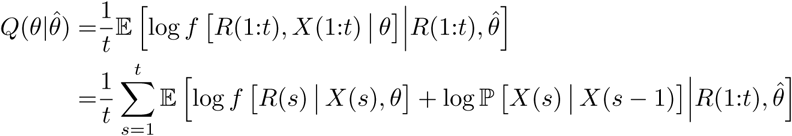

A *maximization* step searches for a maximum of 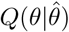 over *θ*. The maximum is then used as the estimate in the next iteration. The crux of the EM method is that 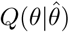 is easier to maximize than the log-likelihood ℓ(*θ*) of the observed data and that improvement in 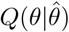 guarantees improvement in ℓ(*θ*).

Mirroring our argument for the Rescorla-Wagner model, we can replace the maximization step with a Newton update:

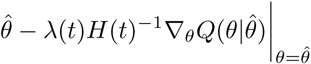

for appropriate learning rate *λ*(*t*) and matrix *H*(*t*). Even in a latent-variable setting, one choice for *H*(*t*) is still the negative of the Fisher’s information matrix scaled be 1*/t* and evaluated at 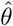, but Newton’s method still requires all rewards up to trial *t*. To recover an online approach, we use a stochastic approximation to 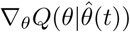 in the maximization step:

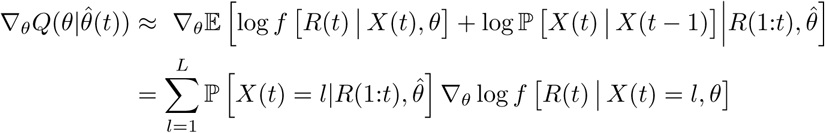

We then replace 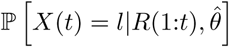, which is computed using all the rewards, with latent-state beliefs *p_l_*(*t*) defined at Eq. 5 which is computed online. This replacement yields the stochastic approximation

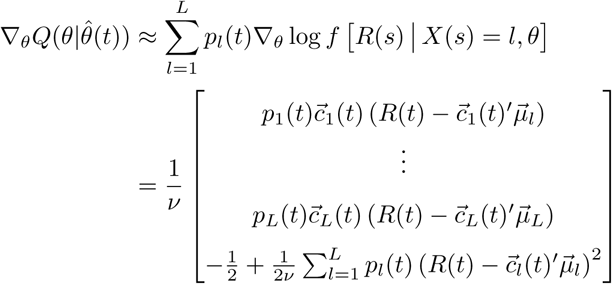

The last equality follows from our assumption that rewards are a Markov chain of Gaussian regression models.

Upon replacing the gradient in Newton’s method with this stochastic approximation, we can arrive at our latent-state learning model if we use *λ*(*t*) = *α*_0_ and matrix

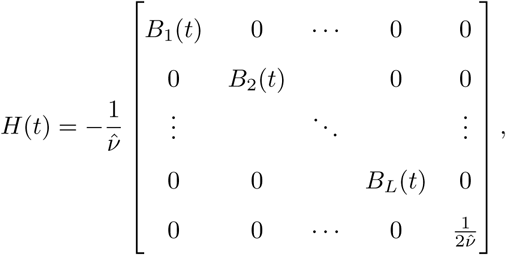

where we used the definition of *B_l_*(*t*) at Eq. 4. For a Markov chain of Gaussian regression models, matrix *H*(*t*) resembles the negative of the Fisher information matrix scaled by 1*/t*:

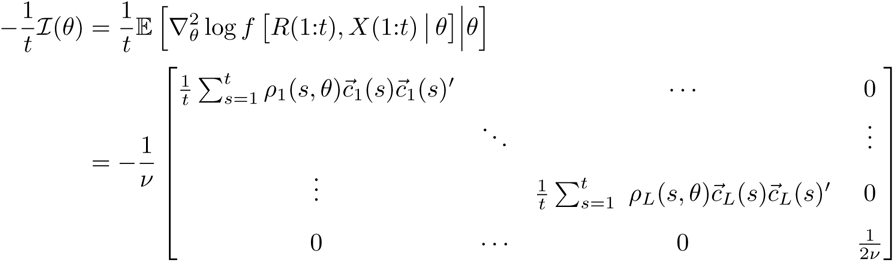

where *ρ_l_*(*s, θ*) = P [*X*(*s*) = *l|θ*]. We simply replaced diagonal blocks with *B_l_*(*t*) which ensures *H*(*t*) is invertible and replaced *ρ_l_*(*s, θ*) with latent state beliefs *p_l_*(*s*). With these choices for *λ*(*t*) and *H*(*t*), we update 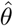 online according to:

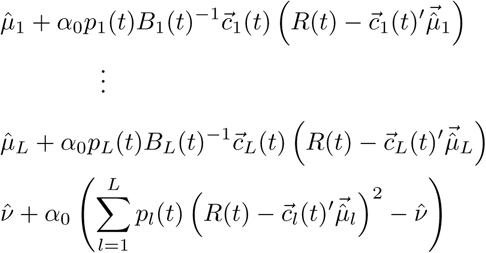

We thus arrive at our latent-state model for updating associative strengths at Eq.(1). We can also recover our latent-state model for updating variance at Eq (7), if we replace the learning rate *α*_0_ with a different learning rate *β*_0_. This completes our motivation for our latent-state model.

## Supporting information

**S1 Text. Additional support and details**. This supplement is divided into four sections: Extensions, Comparison to other learning models, Simulation details, and Sensitivity of predictions to parameters. The first section discusses how to extend the latent-state model to allow for multidimensional rewards and decision-making. The second section examines simulated behavior from other learning models. The third section provides additional details on simulated experiments. The last section examines whether the ability of our model to reproduce target effects in simulation is sensitive to changes in parameters.

**S1 Figure. Hybrid model.** Simulated behavior of the Hybrid model for the same learning experiments in the main text. Gray dashes demarcate experimental stages.

**S2 Figure. Gershman (2017) model with higher concentration parameter.** Simulated behavior of Gershman (2017) model with concentration parameter *α* = 1 for the same learning experiments in the main text. Gray dashes demarcate experimental stages.

**S1 Table. Parameter sensitivity tests.** Quantities measured in each simulation experiment in order to test sensitivity of model predictions to changes in parameters. The value of each test quantity determines whether or not a target learning effect is reproduced by the model.

**S2 Table. Parameter sensitivity results.** Target quantity as a function of target effect (#1–13) and which parameter was varied, keeping all other parameters fixed. By checking whether each target quantity was positive, we could determine whether the model reproduced one of the target effects listed in the main text. In this table, all parameter sets reproduced target effects.

**S3 Table. Parameter sensitivity results *continued.*** Target quantity as a function of target effect (#1–13) and which parameter was varied, keeping all other parameters fixed. By checking whether each target quantity was positive, we could determine whether the model reproduced one of the target effects listed in the main text. In this table, all parameter sets reproduced target effects, except in one case: decreasing parameter *δ* (which controlled unexpected uncertainty) did not reproduce memory modification effect (#13).

## References

1. Huys QJ, Guitart-Masip M, Dolan RJ, Dayan P. Decision-theoretic psychiatry. Clinical Psychological Science. 2015;3(3):400–421.

2. Sutton RS, Barto AG, et al. Reinforcement learning: An introduction. MIT press; 1998.

3. Daw ND, O’doherty JP, Dayan P, Seymour B, Dolan RJ. Cortical substrates for exploratory decisions in humans. Nature. 2006;441(7095):876.

4. Doll BB, Simon DA, Daw ND. The ubiquity of model-based reinforcement learning. Current opinion in neurobiology. 2012;22(6):1075–1081.

5. Wilson R, Collins A. Ten simple rules for the computational modeling of behavioral data. 2019;.

6. Dayan P, Berridge KC. Model-based and model-free Pavlovian reward learning: revaluation, revision, and revelation. Cognitive, Affective, & Behavioral Neuroscience. 2014;14(2):473–492.

7. Gläscher J, Daw N, Dayan P, O’Doherty JP. States versus rewards: dissociable neural prediction error signals underlying model-based and model-free reinforcement learning. Neuron. 2010;66(4):585–595.

8. Rescorla RA, Wagner AR, et al. A theory of Pavlovian conditioning: Variations in the effectiveness of reinforcement and nonreinforcement. Classical conditioning II: Current research and theory. 1972;2:64–99.

9. Pearce JM, Hall G. A model for Pavlovian learning: variations in the effectiveness of conditioned but not of unconditioned stimuli. Psychological review. 1980;87(6):532.

10. Le Pelley ME. The role of associative history in models of associative learning: A selective review and a hybrid model. The Quarterly Journal of Experimental Psychology Section B. 2004;57(3b):193–243.

11. Redish AD, Jensen S, Johnson A, Kurth-Nelson Z. Reconciling reinforcement learning models with behavioral extinction and renewal: implications for addiction, relapse, and problem gambling. Psychological review. 2007;114(3):784.

12. Bouton ME. Context and behavioral processes in extinction. Learning & memory. 2004;11(5):485–494.

13. Vervliet B, Craske MG, Hermans D. Fear extinction and relapse: state of the art. Annual review of clinical psychology. 2013;9:215–248.

14. Redish AD. Addiction as a computational process gone awry. Science. 2004;306(5703):1944–1947.

15. Gershman SJ, Blei DM, Niv Y. Context, learning, and extinction. Psychological review. 2010;117(1):197.

16. Gershman SJ, Hartley CA. Individual differences in learning predict the return of fear. Learning & behavior. 2015;43(3):243–250.

17. Gershman SJ, Niv Y. Exploring a latent cause theory of classical conditioning. Learning & behavior. 2012;40(3):255–268.

18. Gershman SJ, Norman KA, Niv Y. Discovering latent causes in reinforcement learning. Current Opinion in Behavioral Sciences. 2015;5:43–50.

19. Gershman SJ, Monfils MH, Norman KA, Niv Y. The computational nature of memory modification. Elife. 2017;6:e23763.

20. Gershman SJ, Jones CE, Norman KA, Monfils MH, Niv Y. Gradual extinction prevents the return of fear: implications for the discovery of state. Frontiers in behavioral neuroscience. 2013;7:164.

21. Tolman EC. Cognitive maps in rats and men. Psychological review. 1948;55(4):189.

22. Stalnaker TA, Cooch NK, Schoenbaum G. What the orbitofrontal cortex does not do. Nature neuroscience. 2015;18(5):620.

23. Wikenheiser AM, Schoenbaum G. Over the river, through the woods: cognitive maps in the hippocampus and orbitofrontal cortex. Nature Reviews Neuroscience. 2016;17(8):513.

24. Hall G, Pearce JM. Latent inhibition of a CS during CS–US pairings. Journal of Experimental Psychology: Animal Behavior Processes. 1979;5(1):31.

25. Agrawal S, Goyal N. Thompson sampling for contextual bandits with linear payoffs. In: International Conference on Machine Learning; 2013. p. 127–135.

26. Angela JY, Dayan P. Uncertainty, neuromodulation, and attention. Neuron. 2005;46(4):681–692.

27. Wilson RC, Niv Y. Inferring relevance in a changing world. Frontiers in human neuroscience. 2012;5:189.

28. Cappé O. Online EM algorithm for hidden Markov models. Journal of Computational and Graphical Statistics. 2011;20(3):728–749.

29. Page ES. Continuous inspection schemes. Biometrika. 1954;41(1/2):100–115.

30. Granjon P. The CuSum algorithm-a small review. 2013;.

31. Mackintosh NJ. A theory of attention: variations in the associability of stimuli with reinforcement. Psychological review. 1975;82(4):276.

32. Chan SC, Niv Y, Norman KA. A probability distribution over latent causes, in the orbitofrontal cortex. Journal of Neuroscience. 2016;36(30):7817–7828.

33. Rich EL, Wallis JD. Decoding subjective decisions from orbitofrontal cortex. Nature neuroscience. 2016;19(7):973.

34. Niv Y, Daniel R, Geana A, Gershman SJ, Leong YC, Radulescu A, et al. Reinforcement learning in multidimensional environments relies on attention mechanisms. Journal of Neuroscience. 2015;35(21):8145–8157.

35. Kamin LJ. Predictability, surprise, attention, and conditioning. 1967;.

36. Lattal KM, Nakajima S. Overexpectation in appetitive Pavlovian and instrumental conditioning. Animal Learning & Behavior. 1998;26(3):351–360.

37. Rescorla RA. Pavlovian conditioned inhibition. Psychological Bulletin. 1969;72(2):77.

38. Wilson PN, Boumphrey P, Pearce JM. Restoration of the orienting response to a light by a change in its predictive accuracy. The Quarterly Journal of Experimental Psychology Section B. 1992;44(1b):17–36.

39. Rescorla RA. Associative changes in excitors and inhibitors differ when they are conditioned in compound. Journal of Experimental Psychology: Animal Behavior Processes. 2000;26(4):428.

40. Capaldi E. The effect of different amounts of alternating partial reinforcement on resistance to extinction. The American Journal of Psychology. 1957;.

41. Jenkins HM. Resistance to extinction when partial reinforcement is followed by regular reinforcement. Journal of Experimental Psychology. 1962;64(5):441.

42. Theios J. The partial reinforcement effect sustained through blocks of continuous reinforcement. Journal of Experimental Psychology. 1962;64(1):1.

43. Miller RR, Matute H. Biological significance in forward and backward blocking: Resolution of a discrepancy between animal conditioning and human causal judgment. Journal of Experimental Psychology: General. 1996;125(4):370.

44. Monfils MH, Cowansage KK, Klann E, LeDoux JE. Extinction-reconsolidation boundaries: key to persistent attenuation of fear memories. science. 2009;324(5929):951–955.

45. Schiller D, Monfils MH, Raio CM, Johnson DC, LeDoux JE, Phelps EA. Preventing the return of fear in humans using reconsolidation update mechanisms. Nature. 2010;463(7277):49.

46. Ricker ST, Bouton ME. Reacquisition following extinction in appetitive conditioning. Animal Learning & Behavior. 1996;24(4):423–436.

47. Bouton ME, King DA. Contextual control of the extinction of conditioned fear: tests for the associative value of the context. Journal of Experimental Psychology: Animal Behavior Processes. 1983;9(3):248.

48. Brooks DC, Bouton ME. A retrieval cue for extinction attenuates spontaneous recovery. Journal of Experimental Psychology: Animal Behavior Processes. 1993;19(1):77.

49. Li J, Schiller D, Schoenbaum G, Phelps EA, Daw ND. Differential roles of human striatum and amygdala in associative learning. Nature neuroscience. 2011;14(10):1250.

50. Cappé O, Moulines E. On-line expectation–maximization algorithm for latent data models. Journal of the Royal Statistical Society: Series B (Statistical Methodology). 2009;71(3):593–613.

51. Burgess N, Becker S, King JA, O’Keefe J. Memory for events and their spatial context: models and experiments. Philosophical Transactions of the Royal Society of London Series B: Biological Sciences. 2001;356(1413):1493–1503.

52. Gershman SJ, Horvitz EJ, Tenenbaum JB. Computational rationality: A converging paradigm for intelligence in brains, minds, and machines. Science. 2015;349(6245):273–278.

53. Robbins H, Monro S. A stochastic approximation method. In: Herbert Robbins Selected Papers. Springer; 1985. p. 102–109.

54. Dempster AP, Laird NM, Rubin DB. Maximum likelihood from incomplete data via the EM algorithm. Journal of the royal statistical society Series B (methodological). 1977; p. 1–38.

