## Supplementary material for "A flexible and generalizable model of online latent-state learning": S1 Text

#### Extensions

There are two extensions of the model to consider. To extend to multidimensional reward setting, we suppose that the learning agent observes $D$ rewards on each trial: $R_1(t), \ldots, R_D(t)$. In this case, we assume the learning agent updates a separate RW model for each learning state and *each reward dimension*. We replace the cue vector, associative strength, prediction error, associability matrix, and effort matrix with corresponding vectors/matrices $\vec{c}_{l,d}(t), \vec{V}_{l,d}(t), E_{l,d}(t), A_{l,d}(t), B_{l,d}(t)$ that depends on each latent state $l$, trial $t$, and dimension $d$. We update these variables as in the one-dimensional setting with $R(t)$ replaced with the corresponding reward $R_d(t)$. Meanwhile, latent-state beliefs $p_l(t)$ remain specific to a latent state and not the reward dimension. Assuming rewards are independent across dimensions, we replace

$$\phi\left(\frac{E_l(t)}{\sigma(t)}\right),$$

with

$$\prod_d \phi\left(\frac{E_l(t)}{\sigma(t)}\right).$$

We continue to pool variances $\sigma^2(t)$ into a single quantity and thus replace $\overline{E^2}(t)$ in the main text with

$$\overline{E^2}(t) = \frac{1}{D}\sum_{d=1}^{D}\sum_{l=1}^{L} p_l(t) E_{l,d}^2.$$

As before, we pool variances to reduce the number of variables, but one may want to consider separate variances for each latent state and each reward dimension.

Many learning tasks require an agent to decide between $M$ mutually-exclusive options on each trial with rewards generated depending on the agent's decision. To apply our model to such a decision-making environment, we define $\vec{c}_{l,d,m}(t)$ to be the feature vector *if* the agent were to choose option $m$. Then based on the agent's current model on trial $t$, expected reward along dimension $d$ for choosing option $m$ would be

$$W_{d,m}(t) = \sum_{l=1}^{L} p_l(t)\vec{c}_{l,d,m}(t)'V_{l,d}(t)$$

The agent then chooses option m on trial $t$ with probability

$$\frac{\exp \sum_{d=1}^{D} \tau_d W_{d,m}(t)}{\exp\left(\sum_{d=1}^{D}\sum_{n=1}^{M} \tau_d W_{d,n}(t)\right)}$$

for parameter $\tau_d$ determining how much an agent explores versus exploits and weights rewards along each dimension. Note that since the expected reward per option $W_{d,m}(t)$ is available, rules other than the above soft-max rule, e.g. $\epsilon$-greedy rules, could be used to model how an agent makes a decision.

### Comparison to other learning models

We use similar notation to the main text with a vector $\vec{V}(t)$ to denote associative strengths at the start of trial $t$, a vector $\vec{c}(t)$ to denote which cues are presented on trial $t$, $R(t)$ to denote rewards on trial $t$, and $E(t)$ to denote prediction error on trial $t$:

$$E(t) = R(t) - \vec{c}(t)'\vec{V}(t).$$

In addition, we use $C(t)$ to denote the diagonal matrix whose diagonal entries make up the cue vector $\vec{c}(t)$. We assume rewards are one-dimensional unless otherwise specified. Models are extended to decision-making tasks as was done with our latent-state model: estimating the expected reward *for* choosing each option, incorporating these rewards into a softmax function, and choosing an option according to the probability recovered from the softmax function.

In the main text, we compared our latent-state model to the infinite-mixture model in [1] and the Gershman (2017) model in [2]. In prior work, these models helped to explain learning effects not captured by traditional learning models, such as extinction with renewal and memory consolidation [1–3]. Our latent-state model shares a similar Bayesian hierarchical framework to these models whereby rewards are assumed to be constructed in a generative way: a latent state is first randomly drawn on each trial and then rewards are drawn randomly with a distribution that depends on which latent state is drawn. This framework allows learning agents to learn about the hidden structure of their environment. Both models assume a prior for latent states arising from the Chinese Restaurant Process or time-sensitive Chinese Restaurant Process. In contrast, our model assumes latent states evolve according to a discrete-time Markov Chain.

Two quantities are compared in the main text between all models of latent-state learning. First, we compared beliefs in latent state $l$ on trial $t$, which we defined to be the estimated posterior probability of latent state $l$ given the history of cues and rewards. Models of latent-state learning use different approaches to estimate this quantity: the infinite-mixture model uses a particle method, the Gershman (2017) model uses maximum a posteriori (MAP) estimates of latent causes, and our latent-state model uses an approximate Bayesian filter. Second, we compared associative strength of a cue $c$, which we defined to be the estimate of the expected reward if cue $c$ were presented alone where we use $\vec{e}_c$ to denote the cue vector if cue c was presented alone (i.e. a standard basis vector). For our model, belief in latent state $l$ is $p_l(t)$ which estimates the posterior probability of latent state $l$:

$$\mathbb{P}\left[X(t) = l \middle| R(1{:}t), \vec{c}(1{:}t)\right].$$

Associative strength of cue c is $\sum_l V_l(t) p_l(t-1)$ which estimates the expectation

$$\begin{aligned}
&\mathbb{E}\left[R(t) \middle| \vec{c}(t) = \vec{e}_c, R(1{:}t-1), \vec{c}(1{:}t-1)\right] \\
&\quad = \sum_l \mathbb{E}\left[R(t) \middle| X(t) = l, \vec{c}(t) = \vec{e}_c, R(1{:}t-1), \vec{c}(1{:}t-1)\right] \mathbb{P}\left[X(t) = l \middle| \vec{c}(t) = \vec{e}_c, R(1{:}t-1), \vec{c}(1{:}t-1)\right]
\end{aligned}$$

To arrive at expressions of beliefs and associative strengths for the infinite-mixture model, we note that the infinite-mixture model is defined according to Bayes law in Equation (2) from [1] which in our notation is given by:

$$\mathbb{P}\left[X(1{:}t) \middle| R(1{:}t), \vec{c}(1{:}t)\right] \propto \mathbb{P}\left[R(1{:}t), \vec{c}(1{:}t) \middle| X(1{:}t)\right] \mathbb{P}\left[X(1{:}t)\right].$$

In addition, Equation (1) from [1] defines how latent states evolve over time:

$$\mathbb{P}\left[X(t) = k \middle| X(1{:}t-1)\right] = \begin{cases} \frac{a}{t-1+a} & \text{if } k \text{ is a new cause} \\ \frac{N_k}{t-1+a} & \text{otherwise} \end{cases}$$

where $N_k$ are the number of prior latent causes $X(1{:}t-1)$ that are equal to $k$. They then use particle methods to approximate these quantities in a recursive manner. Using these quantities and the law of total probability, they can estimate expected reward given the current cue vector in Equation (3) from [1]. We can thus use Equation (3) from [1] to calculate associative strength of cue c, i.e. the estimated expected reward if cue c were presented alone, by replacing the current cue vector in Equation (3) with $\vec{e}_c$:

$$\begin{aligned}\sum_{X(1:t)} \mathbb{E}\left[R(t)\middle|X(1{:}t), \vec{c}(t) = \vec{e}_c, R(1{:}t-1), \vec{c}(1{:}t-1)\right] \\ \times \mathbb{P}\left[X(t)\middle|\vec{c}(t) = \vec{e}_c, R(1{:}t-1), \vec{c}(1{:}t-1), X(1{:}t-1)\right] \\ \times \mathbb{P}\left[X(1{:}t-1)\middle|R(1{:}t-1), \vec{c}(1{:}t-1)\right].\end{aligned}$$

The middle term can then be expressed by Equation (4) from [1] upon replacing the current cue vector with $\vec{e}_c$:

$$\begin{aligned}&\mathbb{P}\left[X(t)\middle|\vec{c}(t) = \vec{e}_c, R(1{:}t-1), \vec{c}(1{:}t-1), X(1{:}t-1)\right] \\ &\propto \mathbb{P}\left[\vec{c}(t) = \vec{e}_c\middle|X(t), R(1{:}t-1), \vec{c}(1{:}t-1), X(1{:}t-1)\right] \mathbb{P}\left[X(t)\middle|X(1{:}t-1)\right].\end{aligned}$$

Lastly, beliefs in latent state $l$ for the infinite mixture model is the posterior distribution of latent states given observations:

$$\mathbb{P}\left[X(t) = l\middle|R(1{:}t), \vec{c}(1{:}t), X(1{:}t-1)\right].$$

To arrive at an expression of beliefs and associative strengths for the Gershman (2017) model, we note that Equation (9) in [2] denotes the expected reward given the current cue vector and past information. Thus, replacing the current cue vector with $\vec{e}_c$ gives the following expression:

$$E\left[R(t)\middle|\vec{c}(t) = \vec{e}_c, R(1{:}t-1), \vec{c}(1{:}t-1)\right] = \sum_l w_{lc}\mathbb{P}\left[X_t = l\middle|\vec{c}(t) = \vec{e}_c, R(1{:}t-1), \vec{c}(1{:}t-1), W^n\right]$$

where $W^n$ collects the estimated associative weights in the Gershman (2017) model and $w_{lc}$ denotes the associative weight for cue $c$ and latent state $l$. Meanwhile, Equation (8) in [2] gives an expression for posterior distribution of latent states using observations up to and including trial $t$:

$$\mathbb{P}\left[X(t) = l\middle|R(1{:}t), \vec{c}(1{:}t), W^n\right].$$

Thus, the estimate to this expression defines beliefs in latent state $l$ for the Gershman (2017) model. Beliefs and associative strengths were estimated using associated code with the infinite-mixture model and with the Gershman (2017) model, modified to estimate associative strengths and to allow for decision-making.

We also considered three models of traditional associative learning: the RW model and two additional models. We considered the feature reinforcement learning with decay (fRL+decay) model in [4], since this model provided the best explanation of behavioral and neural data of the 3-arm bandit task in [4], one of the experiments simulated in the current paper. The fRL+decay model updates associative strength in a similar manner to the RW model for cues that are present, but otherwise decays associative strength to zero for cues that are absent to allow for 'forgetting' of the associative strength. That way, the model captures how one might forget over time about associations between cues and rewards. These updates can be written concisely as:

$$\vec{V}(t+1) = \vec{V}(t) + \kappa_{fRL}\vec{c}(t)E(t) - \eta_{fRL}(I - C(t))\vec{V}(t)$$

for a learning rate parameter $\kappa_{fRL}$ and a decay rate parameter $\eta_{fRL}$. We set parameters $\kappa_{fRL} = 0.15$ and $\eta_{fRL} = 0.45$ to approximately match parameter estimates in Table 1 of [4].

We also considered the Hybrid model in [5], since this model allows associability to vary in time. This model updates associative strength in a similar manner to the RW model:

$$\vec{V}(t+1) = \vec{V}(t) + \kappa_H A(t)\vec{c}(t)E(t)$$

where $\kappa_H$ is a constant learning rate parameter and $A(t)$ is an matrix capturing associability. The benefit of the Hybrid model, however, is that associability $A(t)$ is updated based on prediction errors. Specifically, associability $A(t)$ is a diagonal matrix that is initialized to be the identity matrix and updated according to

$$A(t+1) = A(t) + \eta_H C(t)\left(A(t) - |E_d(t)|C(t)\right)$$

where $\eta_H$ is a constant parameter. Hence, associability increases when the model poorly predicts rewards, allowing for quicker learning in uncertain environments. In all simulation experiments, we fixed parameters $\eta_H = 0.857$ and $\kappa_H = 0.3$ and initialized each associative strength entry in $\vec{V}_d(0)$ to zero and each associability entry in the diagonal of $A(0)$ to 0.926. Except for $\kappa_H$, these parameters are found in the Supplement of [5]. The supplement of [5] uses a value of 0.166 for $\kappa_H$, which we found poorly highlighted the associability process in simulation experiments. We thus increased $\kappa_H$. The simulated examples demonstrate the Hybrid model with the given parameters captures some, but not all of the learning effects (Fig S1). For example, the Hybrid captured blocking, overexpectation, and conditioned inhibition, but did not capture backwards blocking or the experiments from Rescorla [6] and Wilson et al. [7]. The model also did not capture the partial reinforcement extinction effect [8–10] and the rapid return of expectations during renewal [11].

We also considered another model of latent-state learning: a model from Redish et al [12]. This model combines two processes: state classification and value learning. State classification is performed using radial basis functions to classify each trial into one of a number of states based on the combination of cues and rewards observed on that trial. Value learning uses the classification of states to perform a temporal-difference learning update to estimate the state-action value function of the optimal policy, i.e. the expected return for choosing a particular action in a particular state and then following the optimal policy. In order to compare the Redish (2007) model to other models of latent-state learning, we defined belief in a latent state to be 1 if the state was identified as the current agent state and 0 otherwise. Associative strength for each cue was defined as the average estimated value between reinforcing and not reinforcing the cue presented alone. We use this definition rather the expected reward given a cue, since the Redish (2007) model places value only on state-action pairs and does not generate expectations. Further, states are only classified if rewards are known in addition to cues. Parameters were taken directly from the code accompanying Redish et al [12]. Further, cues and rewards were scaled by 100 as was done in the code that accompanies the paper in [12].

Lastly, we again considered the Gershman (2017) model but with a higher concentration parameter ($\alpha = 1$). We used a larger concentration parameter, because we found the smaller $\alpha = 0.1$ relied on one latent state for a number of simulation experiments (i.e. beliefs for latent state 1 were near one). For both concentration parameters, the Gershman (2017) model could capture blocking, overexpectation, and conditioned inhibition, but could not capture the experiments from Rescorla [6] or Wilson et al [7] (Fig S2). Both concentration parameters also led to similar predictions for associative strengths for partial reinforcement effect, renewal, and spontaneous recovery experiments, failing to capture the target effect in each experiment. Unlike the lower concentration parameter, the higher concentration parameter, however, did not lead to a reconsolidation window, i.e. a set of time lags between retrieval and extinction in which the associative strength of the cue could be decreased during testing.

#### Simulation details

For most experiments, we represented cue presentation on trial $t$ as vectors $\vec{c}(t)$ such that each entry in the vector corresponded to a particular cue (e.g., cue A) and took a value of 1 if the cue was present and 0 otherwise. We encoded binary rewards as -1/2 or 1/2 for our latent-state model, and so, associative strengths are often shifted by 1/2 for our model when compared to other models.

#### Blocking, overexpectation, and conditioned inhibition

For these experiments, we followed conditioning schedules in [13]. Blocking simulations consisted of 20 trials of reinforcing Cue A, followed by 16 trials alternating between reinforcing a compound of Cue A and Cue B and reinforcing a compound of Cue C and Cue D. Overexpectation simulations consisted of 100 trials alternating between reinforcing Cue A and reinforcing Cue B, followed by 10 trials of reinforcing a compound of Cue A and Cue B. Conditioned inhibition simulations consisted of 200 trials alternating between reinforcing Cue A and weakly reinforcing (i.e. $R(t) = 1/2$) a compound of Cue A and Cue B.

#### Backwards blocking

We used a similar conditioning schedule of the blocking experiment: 20 trials of reinforcing Compound AB, followed by 16 trials of reinforcing Cue A.

#### Experiments 1A and 1B from Rescorla (2000) [6]

We followed the conditioning schedule in the original experiment [6]. For both experiments, the first stage of training involved presenting the five trial types (A+,C+,X+,BX,DX) in random order for a total of 500 trials ($= [8 \times 12$ days $+ 4 \times 1$ day$] \times 5$ trial types). In Stage 2, experiment 1A followed this conditioning by presenting the reinforced compound AB+ for a total of 8 times, whereas experiment 1B followed this conditioning by presenting the non-reinforced compound AB 8 times. We measured the change in associative strength from the end of Stage 1 to the end of Stage 2.

#### Experiment 1 from Wilson et al (1992) [7]

We followed the conditioning schedule used to simulate this experiment in the latent-state modeling paper in [1]. For Group C, this schedule consisted of 50 trials of alternating between reinforcing a light and tone with food and presenting a light and tone without food, followed by 10 trials of reinforcing the light with food.

#### Partial reinforcement extinction effect (PREE)

For Experiment 1 of PREE, we followed the conditioning schedule used to simulate PREE in the latent-state modeling paper in [1]. For the partial reinforcement group, the schedule consisted of 20 trials of partial reinforcement (alternating between reinforcing and not reinforcing a trial) followed by 20 trials of no reinforcement. For the continuous reinforcement group, the schedule consisted of 20 trials of continuous reinforcement followed by 20 trials of no reinforcement. Experiment 2 of PREE was identical for Experiment 1 except that 20 trials of continuous reinforcement was added for both groups between the initial 20 trials of reinforcement and the final 20 trials of extinction.

##### Renewal and spontaneous recovery

In these experiments, we started with a similar schedule to our PREE simulation experiments: 20 trials of continuous reinforcement (acquisition), followed by 20 trials of no reinforcement (extinction). These stages were then followed by 10 trials of reinforcement (renewal). For the renewal experiment, two experimental conditions are considered in which either the same context is provided throughout each of the three phases (AAA) or the same visual/spatial context is provided during acquisition and renewal phases with a new context provided during the extinction phase (ABA). For all models except our latent-state model, cue vectors encoded context. For our latent-state model, context was captured by re-setting beliefs to be split equally among latent states when context shifts. This was used to capture the ability of a context to signal to the participant that need to be open to starting a new model in the presence of new cue or return to an old model in the presence of an old cue. For the spontaneous recovery experiment, we added a shift in temporal context by delaying renewal by 200 units after extinction.

##### Memory modification

We followed the simulation experiment in [2], which was designed to reflect the experiments in Monfils et al [14] and Schiller et al [15]. The experiment consisted of four phases. First a cue was reinforced on three trials for an acquisition phase. Second, the cue is presented once but not reinforced for a 'retrieval' phase. Third, the cue is presented for 18 trials but not reinforced for a 'extinction' phase, and fourth, the cue is presented and not reinforced for a final 'test' phase. Further, we assumed that the time (with same units as inter-trial time) between each phase was 0, 20, x, and 200, respectively. Parameter x represents the time between retrieval and extinction and was varied from 0 to 150. The Gershman (2017) model was simulated as was done in [2].

#### Sensitivity to parameters

We evaluated sensitivity of model predictions to changes in parameters. To be able to examine sensitivity, we needed quantities upon which to measure sensitivity. For each simulated experiment, we thus identified a test quantity whose value determined whether a target effect (see Table 1 from main text) was reproduced by the model or not. When positive, each test quantity implied the target effect was reproduced by the model. For example, overexpectation was evaluated based on whether the associative strength of cue A decreased when reinforced together with cue B after each cue had initially been reinforced separately. We thus can determine if our model reproduces this effect by measuring the difference in associative strength of cue A after being reinforced separately (i.e. at the end of stage 1) compared to its associative strength after being reinforced as a compound (i.e. at the end of stage 2). If this quantity is greater than zero, then indeed the associative strength decreased as desired. Target quantities we proposed are listed for each simulated experiment in Table S1.

With 13 test quantities defined, we varied each parameter by $\pm 10\%$ of the fixed value used for all simulation experiments in the main text. All experiments were simulated for each new value, keeping the other parameters fixed. In each case, we measured the test quantity and check whether its value was positive, which would determine if the model reproduced the target effect for the given set of parameters. The results are summarized in Tables S2–S3. All parameter sets reproduced target effects, except in one case: decreasing parameter $\delta$ (which controlled unexpected uncertainty) did not reproduce memory modification effect (#13).

#### References


1. Gershman SJ, Niv Y. Exploring a latent cause theory of classical conditioning. Learning & behavior. 2012;40(3):255–268.

2. Gershman SJ, Monfils MH, Norman KA, Niv Y. The computational nature of memory modification. Elife. 2017;6:e23763.

3. Gershman SJ, Blei DM, Niv Y. Context, learning, and extinction. Psychological review. 2010;117(1):197.

4. Niv Y, Daniel R, Geana A, Gershman SJ, Leong YC, Radulescu A, et al. Reinforcement learning in multidimensional environments relies on attention mechanisms. Journal of Neuroscience. 2015;35(21):8145–8157.

5. Li J, Schiller D, Schoenbaum G, Phelps EA, Daw ND. Differential roles of human striatum and amygdala in associative learning. Nature neuroscience. 2011;14(10):1250.

6. Rescorla RA. Associative changes in excitors and inhibitors differ when they are conditioned in compound. Journal of Experimental Psychology: Animal Behavior Processes. 2000;26(4):428.

7. Wilson PN, Boumphrey P, Pearce JM. Restoration of the orienting response to a light by a change in its predictive accuracy. The Quarterly Journal of Experimental Psychology Section B. 1992;44(1b):17–36.

8. Capaldi E. The effect of different amounts of alternating partial reinforcement on resistance to extinction. The American Journal of Psychology. 1957;.

9. Theios J. The partial reinforcement effect sustained through blocks of continuous reinforcement. Journal of Experimental Psychology. 1962;64(1):1.

10. Jenkins HM. Resistance to extinction when partial reinforcement is followed by regular reinforcement. Journal of Experimental Psychology. 1962;64(5):441.

11. Bouton ME. Context and behavioral processes in extinction. Learning & memory. 2004;11(5):485–494.

12. Redish AD, Jensen S, Johnson A, Kurth-Nelson Z. Reconciling reinforcement learning models with behavioral extinction and renewal: implications for addiction, relapse, and problem gambling. Psychological review. 2007;114(3):784.

13. Le Pelley ME. The role of associative history in models of associative learning: A selective review and a hybrid model. The Quarterly Journal of Experimental Psychology Section B. 2004;57(3b):193–243.

14. Monfils MH, Cowansage KK, Klann E, LeDoux JE. Extinction-reconsolidation boundaries: key to persistent attenuation of fear memories. science. 2009;324(5929):951–955.

15. Schiller D, Monfils MH, Raio CM, Johnson DC, LeDoux JE, Phelps EA. Preventing the return of fear in humans using reconsolidation update mechanisms. Nature. 2010;463(7277):49.
