## Supplementary material for "A flexible and generalizable model of online latent-state learning": S1 Table

| # | Experiment | Test quantity |
|---|---|---|
| (1) | Blocking | Difference in associative strength of cue C vs cue B at task end |
| (2) | Overexpectation | Difference in associative strength of cue A at stage 1 end vs stage 2 end |
| (3) | Conditioned inhibition | Negative of associative strength of cue B at task end |
| (4) | Backwards blocking | Difference in associative strength of cue B at stage 1 end vs stage 2 end |
| (5) | Rescorla (2000) 1A | Difference in change in associative strength of cue B vs cue A across stage 2 |
| (6) | Rescorla (2000) 1B | Difference in change in associative strength of cue B vs cue A across stage 2 |
| (7) | Wilson et al (1992) 1 | Difference in associative strength of cue A at task end for Group E vs Group C |
| (8) | PREE Exp 1 | Difference in associative strength of cue A in partial reinforcement group vs continuous reinforcement group at start of stage 2 (i.e. extinction) |
| (9) | PREE Exp 2 | Difference in associative strength of cue A in partial reinforcement group vs continuous reinforcement group at start of stage 3 (i.e. extinction) |
| (10) | Renewal (rapid return) | Difference in expected rewards on second trial of stage 3 (renewal) vs second trial of stage 1 (acquisition) |
| (11) | Renewal (w/ context) | Difference in expected rewards on second trial of stage 3 (renewal) with visual/spatial context shift vs no context shift |
| (12) | Spontaneous recovery | Difference in associative strength of a cue on second trial of stage 3 (renewal) with temporal context shift vs no temporal context shift |
| (13) | Memory modification | Difference in associative strength of a cue on test trial with time delay of 5 after retrieval vs time delay of 1 |

**Table S1.** Quantities measured in each simulation experiment in order to test sensitivity of model predictions to changes in parameters. The value of each test quantity determines whether or not a target learning effect is reproduced by the model.
