## Supplementary material for "A flexible and generalizable model of online latent-state learning": S2 Table

| # | $\alpha_0$ 0.045 | $\alpha_0$ 0.055 | $\beta_0$ 0.045 | $\beta_0$ 0.055 | $\gamma$ 0.045 | $\gamma$ 0.055 | $\sigma_0$ 0.45 | $\sigma_0$ 0.55 |
|---|---|---|---|---|---|---|---|---|
| (1) | 0.070 | 0.083 | 0.082 | 0.082 | 0.083 | 0.083 | 0.083 | 0.091 |
| (2) | 0.324 | 0.327 | 0.327 | 0.327 | 0.327 | 0.327 | 0.327 | 0.330 |
| (3) | 0.250 | 0.250 | 0.250 | -0.250 | 0.250 | 0.250 | 0.250 | 0.250 |
| (4) | 0.067 | 0.065 | 0.071 | 0.071 | 0.071 | 0.071 | 0.071 | 0.075 |
| (5) | 1.000 | 1.000 | 1.000 | 1.000 | 1.000 | 1.000 | 1.000 | 1.000 |
| (6) | 1.000 | 1.000 | 1.000 | 1.000 | 1.000 | 1.000 | 1.000 | 1.000 |
| (7) | 0.090 | 0.083 | 0.073 | 0.092 | 0.073 | 0.073 | 0.073 | 0.059 |
| (8) | 0.308 | 0.292 | 0.293 | 0.293 | 0.292 | 0.292 | 0.292 | 0.277 |
| (9) | 0.003 | 0.002 | 0.002 | 0.002 | 0.002 | 0.002 | 0.002 | 0.002 |
| (10) | 0.150 | 0.228 | 0.221 | 0.262 | 0.230 | 0.230 | 0.230 | 0.277 |
| (11) | 0.095 | 0.037 | 0.044 | 0.014 | 0.037 | 0.037 | 0.037 | 0.010 |
| (12) | 0.044 | 0.018 | 0.021 | 0.007 | 0.018 | 0.018 | 0.018 | 0.005 |
| (13) | 0.002 | 0.002 | 0.002 | 0.002 | 0.002 | 0.002 | 0.001 | 0.003 |

**Table S2.** Target quantity as a function of target effect (#1–13) and which parameter was varied, keeping all other parameters fixed. By checking whether each target quantity was positive, we could determine whether the model reproduced one of the target effects listed in the main text. In this table, all parameter sets reproduced target effects.
