## Supplementary figures and images for "A flexible and generalizable model of online latent-state learning"

### S1 Figure

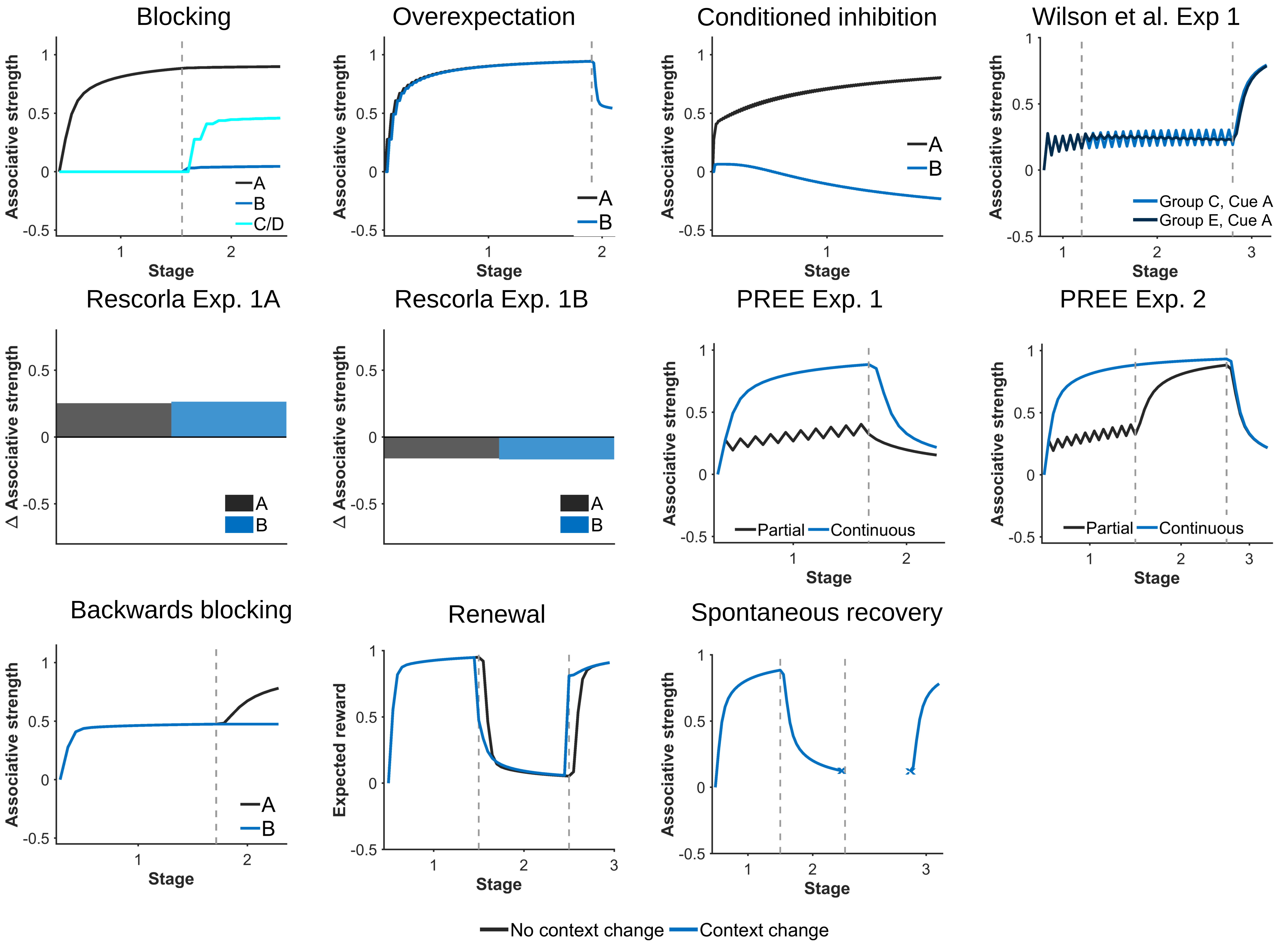

### S2 Figure

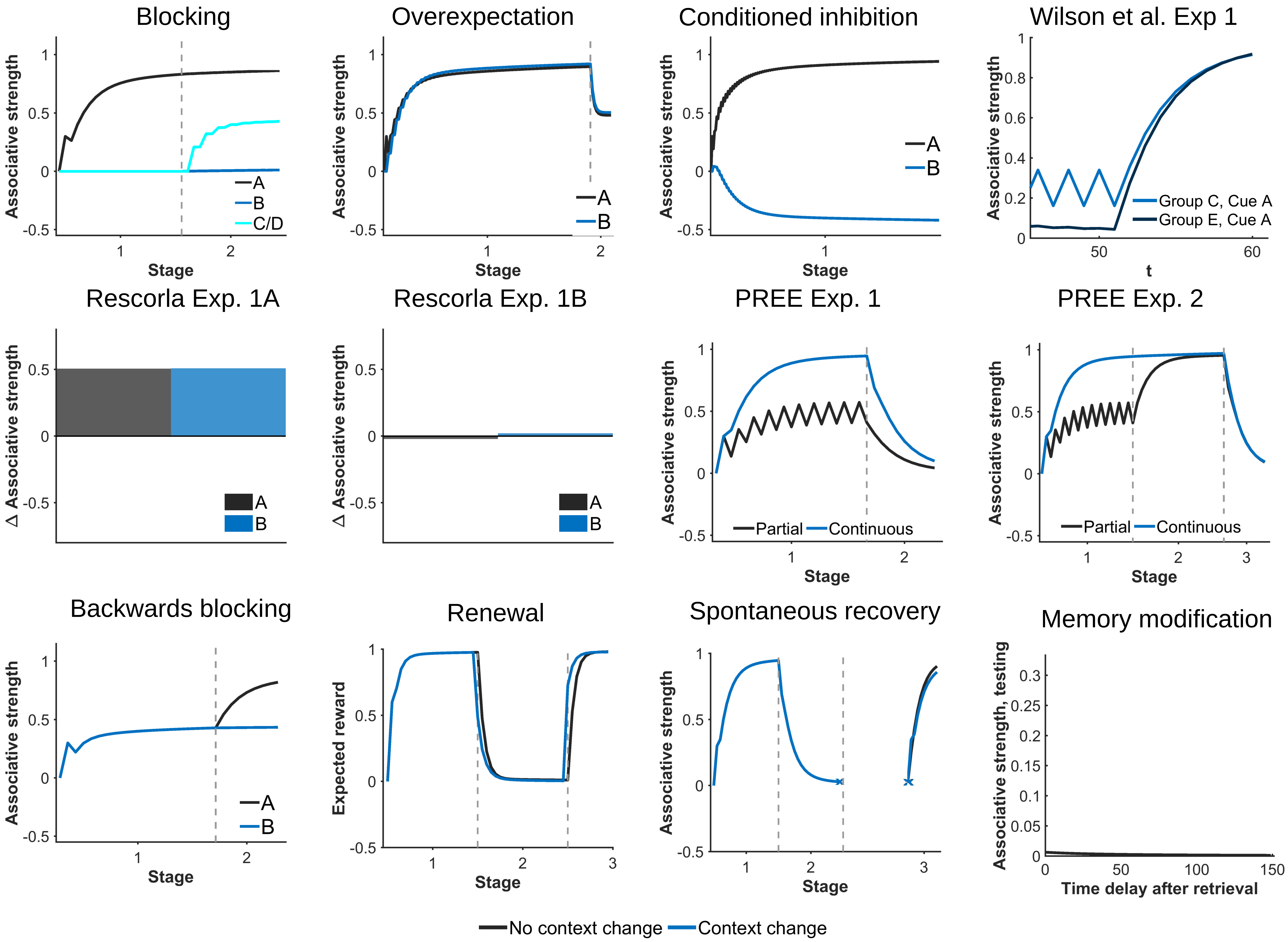
